# The intracellular respiratory burst of plant immunity signaling

**DOI:** 10.64898/2026.11.18.767486

**Authors:** Pedro Barreto, Elias Feitosa-Araujo, Lena Sophie Knorr, Maxim Limbächer, Jan-Ole Niemeier, Melissa Szymlek, Guillaume Decros, Ana Carolina Vilchez, Subhankar Raha, Christiane Rohrbach, Amanta Seifried, Jule Niebisch, Daniela Döben, Ke Zheng, Alina Althaus, Ana Cislaghi, Richard Jaeger, Marcelle Ferreira Silva, Jan Multhoff, Sophie Lichtenauer, Stephan Wagner, Eliza Loo, Adriano Nunes-Nesi, Veronica G. Maurino, Stefanie Müller-Schüssele, Till Ischebeck, Cyril Zipfel, Alisdair R. Fernie, Lars M. Voll, Markus Schwarzländer

## Abstract

Pattern-triggered immunity (PTI) is classically associated with an extracellular respiratory burst generated by plasma membrane-localized NADPH oxidases (RBOHs). Here, we identify an ancient RBOH-independent intracellular respiratory burst driven by central carbon metabolism. Using in vivo biosensors, we demonstrate that receptor activation rapidly triggers sugar catabolism via glycolysis, shifting the NAD pool toward its reduced state. This redox pressure drives a burst in mitochondrial respiratory flux and oxygen consumption. Consequent mitochondrial reactive oxygen species (ROS) production activates ANAC017-mediated retrograde signaling to induce broader PTI gene expression. This “intracellular respiratory burst” represents a conserved early hallmark of immunity across land plants and links cellular carbon status to immune activation. It reveals functional parallels between plant immune metabolism and metabolic reprogramming during mammalian immune-cell activation.

## Main Text

Plants rely on innate immune mechanisms operating at the cellular level to recognize potential pathogen threats and coordinate defense strategies. Surface-localized pattern recognition receptors (PRRs) detect conserved elicitors, initiating pattern-triggered immunity (PTI). This fundamental recognition event triggers a rapid cascade of signaling processes, including calcium (Ca^2+^) influx, mitogen-activated protein kinase (MAPK) activation, broad transcriptomic reprogramming and a major apoplastic burst of reactive oxygen species (ROS) (*1*). The extracellular ROS burst is produced by plasma membrane-localized NADPH oxidases, termed Respiratory Burst Oxidase Homologs (RBOHs), and plays a central role in signal transduction and restricting pathogen entry. Because RBOH activity relies on cellular metabolism to supply reducing equivalents in the form of NADPH, PRR activation inherently imposes a major demand on carbon and redox metabolism (*2–6*).

While major changes in primary metabolism and intracellular ROS dynamics were observed during infection, and specifically PTI, in many instances, these were often linked directly or indirectly to RBOH activity (*2*). Intracellular signatures of H_2_O_2_ dynamics that were initially interpreted as RBOH-derived (*3*) were subsequently demonstrated to be RBOH-independent (*4*). Hexose import through Sugar Transporters (STPs) from the apoplast to the cytosol during PTI was initially interpreted mainly as an apoplast clearance response to remove resources from the reach of the pathogen (*5*, *6*). A central role for mitochondria in plant responses to microbes was suggested more than two decades ago (*7*), partly based on models of ROS generation and cell death developed for mammalian systems. Since then, biotic stress responses were observed to depend on mitochondrial electron transport function and carbon metabolism (*8–10*). Yet, mechanistic insight remained difficult to obtain due to the pleiotropic nature that modifications of central metabolic functions generally entail.

Reprogramming of nuclear gene expression in response to mitochondrial dysfunction in transcriptomic studies of retrograde signaling was noted to share similarities with biotic stress responses (*11*). Yet, while key components and mechanisms of mitochondrial retrograde signaling were subsequently identified (*12*), the link between mitochondrial signaling and plant biotic stress responses remained at the observational level, and models of plant-microbe interaction do not typically include mitochondria. That is despite recent observations that implicate mitochondrial dynamics and intracellular positioning in plant defense responses. Mitochondria immobilize and oxidize at fungal entry sites to coordinate entry resistance (*13*), while MPK3/6-mediated phosphorylation of the mitochondrial motility protein MIRO1 promotes guard-cell mitochondrial fusion to restore membrane potential and organic acid metabolism during stomatal closure (*14*). In a recent study, mitochondrial ROS were shown to stimulate physical mitochondria-ER contact sites to facilitate ROS diffusion into the ER, initiating an unfolded protein response (UPR) and activating stress-related immunity independently of apoplastic ROS (*15*). Despite these and other individual observations, how PRR activation, mitochondrial function, and retrograde nuclear reprogramming may be linked mechanistically during PTI remains undefined.

The work described here addresses this gap by identifying a land-plant-conserved “intracellular respiratory burst” that is a hallmark of a metabolic branch of PTI signaling and acts in parallel to canonical RBOH-dependent extracellular ROS production (Fig. 1A). PRR perception rapidly mobilizes central carbon metabolism, driving glycolytic sugar catabolism and altering cytosolic redox state (NADH/NAD^+^). This sudden metabolic redox pressure increases electron flux through the mitochondrial ETC, generating an intracellular burst in tissue oxygen consumption and driving mitochondrial reactive oxygen species (ROS) production to give rise to sustained intracellular hydrogen peroxide H_2_O_2_ and glutathione redox potential (*E*_GSH_) signatures. Mitochondrial-derived H_2_O_2_ triggers ANAC017-dependent retrograde signaling to induce the mitochondrial dysfunction regulon (MDR) contributing to the transcriptomic defense response and modulating defense outcome.

**Figure 1.**
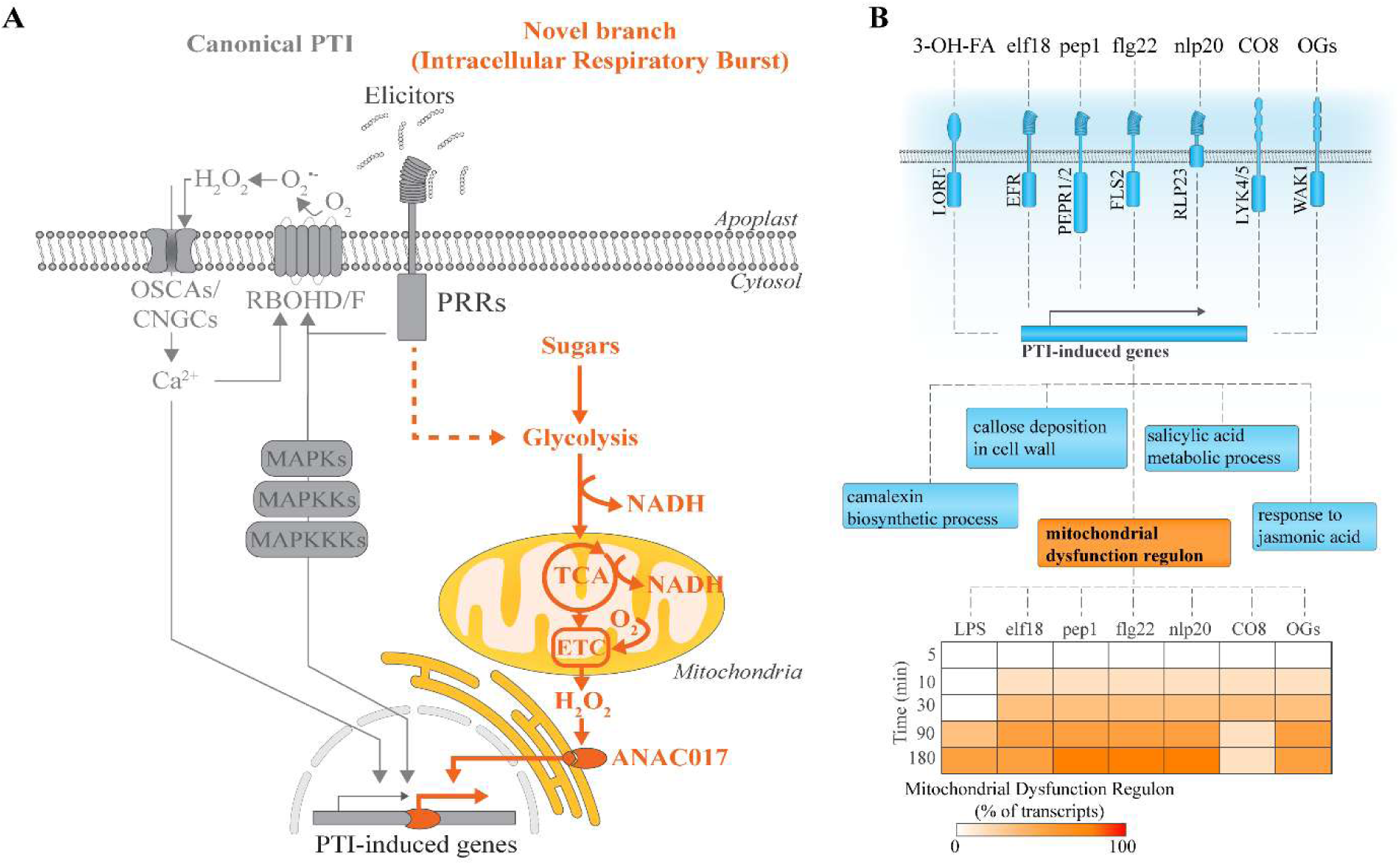
A novel branch of pattern-triggered immunity (PTI). **(A)** Proposed model of the novel metabolic branch of PTI described in this study. An intracellular respiratory burst represents an integral component of PTI that operates in parallel with the canonical PTI branch. This branch involves rapid mobilization of central carbon metabolism, including increased glycolytic flux and respiration, resulting in a shift of the cytosolic NAD redox state towards reduction. Increased electron pressure at the mitochondrial electron transport chain leads to sustained mitochondrial H₂O₂ production, which activates the canonical ANAC017 pathway and induces the expression of mitochondrial dysfunction regulon transcripts. OSCA, REDUCED HYPEROSMOLALITY-INDUCED Ca^2+^ INCREASE; CNGC, CYCLIC NUCLEOTIDE-GATED CHANNEL. **(B)** Mitochondrial dysfunction is a component of PTI. A meta-analysis of the dataset published by Bjornson et al. (2021), in which Arabidopsis seedlings were treated with a range of different elicitors, revealed that the mitochondrial dysfunction regulon (MDR) is among the gene sets enriched in the upregulated transcripts (Table S1). The transcripts commonly induced by all elicitors were classified as PTI-induced genes and subjected to gene enrichment analysis. The response of MDR transcripts to each individual elicitor is shown in Table S2. 3-OH-FA, bacterial hydroxylated fatty acid; LORE, S-lectin receptor kinase LIPOOLIGOSACCHARIDE-SPECIFIC REDUCED ELICITATION; elf18, 18-amino-acid epitope derived from bacterial elongation factor Tu; EFR, LRR-RK EF-TU RECEPTOR; pep1, 23-amino-acid peptide potentially released upon cellular damage; PEPR1/2, LRR-RKs PEP1 RECEPTOR 1 and 2; flg22, 22-amino-acid epitope derived from bacterial flagellin; FLS2, FLAGELLIN SENSING 2; nlp20, 20-amino-acid peptide derived from bacterial, oomycete, and fungal necrosis- and ethylene-inducing peptide 1-like proteins; RLP23, RECEPTOR-LIKE PROTEIN 23; CO8, octameric fragment of fungal cell walls; LYK4/5, LYSM-CONTAINING RECEPTOR KINASE 4 and 5; OGs, oligogalacturonides derived from the plant cell wall; WAK1, WALL-ASSOCIATED KINASE 1.

### The PTI response includes mitochondrial retrograde signaling

Previously, we made the surprising observation that pathogen response-associated transcripts are among the most responsive when mitochondrial function is impaired in Arabidopsis leaves (*11*). Vice versa, re-analysis of a transcriptomic study of the Arabidopsis PTI response to seven elicitors (*16*) revealed the MDR, a set of 24 transcripts that respond to genetic and pharmacological disruption of plant mitochondrial function (*17*), as a common feature (Fig. 1B; Table S1, S2). The transcriptomic response to the mitochondrial complex III inhibitor antimycin A overlapped by >50% with PTI-responsive transcripts (Fig. S1A, B; Table S3). Since a major proportion of mitochondrial retrograde signaling to induce MDR is mediated by NAC-transcription factors, most prominently ANAC017 (*18*), we compared the response of 970 core PTI-responsive transcripts (*16*) to the transcriptomic response to ANAC017 overexpression (*19*) and found a strong overlap (Fig. S1C, D; Table S3). These analyses indicate involvement of mitochondrial (dys-)function and ANAC017-mediated retrograde signaling in PTI.

We next quantified the flg22 response of three MDR transcripts: ALTERNATIVE OXIDASE 1A (*AOX1a*), OUTER MITOCHONDRIAL MEMBRANE PROTEIN OF 66 KDA (*OM66*), and OXIDATIVE SIGNAL-INDUCIBLE1 (*OXI1*), and three other PTI-induced transcripts: *RBOHD,* VACUOLELESS GAMETOPHYTE (*VLG*), and CYTOCHROME P450, FAMILY 81, SUBFAMILY F, POLYPEPTIDE 2 (*CYP81F2*) in wild type and the *fec* (*fls2c efr cerk1*) background, which lacks flg22, elf18 and chitin receptors (Fig. 2A, B). Induction of all transcripts was almost abolished in *fec*, indicating that mitochondrial retrograde signaling is induced downstream of the flg22 receptor FLS2. The MDR response depended critically on ANAC017, since it was lost in two ANAC017 mutant backgrounds, *anac017-1* and *rao2-1* (Fig. 2A, C). *RBOHD* and *CYP81F2* showed a similar dependence, whereas *VLG* did not. While *RBOHD* induction required ANAC017 and *RBOHD* transcript abundance was elevated by ANAC017 overexpression (*19*), the initial extracellular ROS burst was maintained in both *anac017* mutant backgrounds, suggesting that the RBOH-module of PTI signaling remains operational upon impairment of mitochondrial retrograde signaling (Fig. 2D). Moreover, *anac017* plants showed decreased bacterial titers in infection assays using bioluminescent *Pseudomonas syringae* pv. tomato DC3000 (*20*), whereas titers increased in the *fec* control, suggesting a hitherto unrecognized immune-regulatory role of ANAC017-mediated mitochondrial retrograde signaling in PTI (Fig. 2E, F).

**Figure 2.**
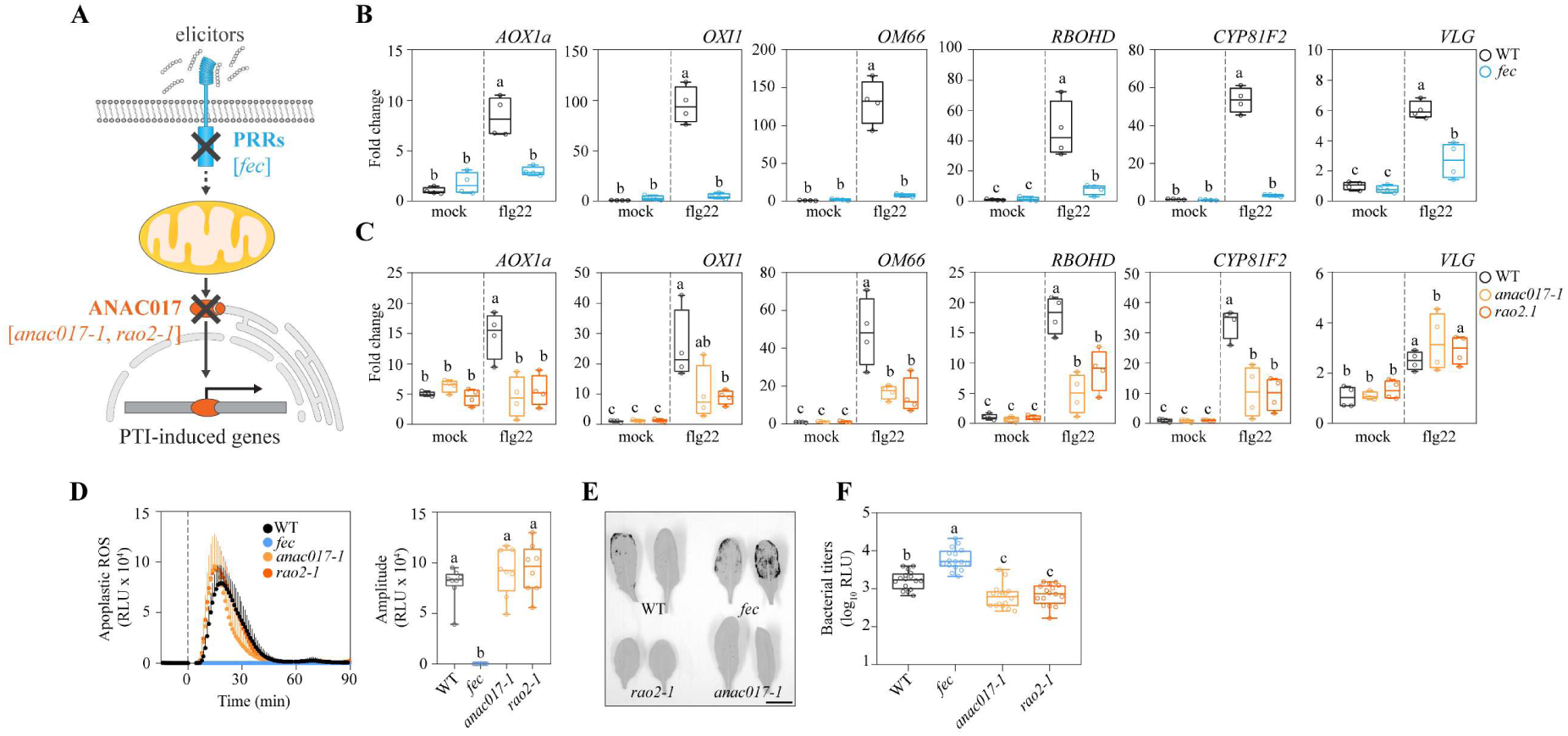
ANAC017-mediated mitochondrial retrograde signaling in PTI. **(A)** Schematic representation of the hypothesized role of ANAC017 in PTI. **(B-C)** Relative transcript abundance of ALTERNATIVE OXIDASE 1A (*AOX1a*), OXIDATIVE SIGNAL-INDUCIBLE 1 (*OXI1*), OUTER MITOCHONDRIAL MEMBRANE PROTEIN OF 66 KDA (*OM66*), RESPIRATORY BURST OXIDASE HOMOLOG D (*RBOHD*), CYTOCHROME P450, FAMILY 81, SUBFAMILY F, POLYPEPTIDE 2 (*CYP81F2*), and VACUOLELESS GAMETOPHYTE (*VLG*) in leaves of 4-week-old *Arabidopsis thaliana* plants. Transcript abundance was quantified by RT-qPCR 50 min after treatment with 100 nM flg22 or water (mock) in **(B)** WT and *fec* (*fls2c efr cerk1*) and **(C)** WT, *anac017-1,* and *rao2-1*. Transcript levels were normalized to WT mock control. n = 4 biological replicates; error bars represent minimum to maximum values. **(D)** Apoplastic ROS burst dynamics in WT, *fec*, *anac017-1,* and *rao2-1* following treatment with 100 nM flg22 (left). Dashed lines indicate the time of flg22 application. The amplitude of the flg22-induced apoplastic ROS burst is shown on the right. n = 8 biological replicates; error bars represent minimum to maximum values. **(E)** Representative chemiluminescent images of leaves from WT, *fec*, *anac017-1*, and *rao2-1* infiltrated with *Pseudomonas syringae* pv. tomato DC3000 carrying a luciferase operon (OD = 0.001). Images are representative of the experiment shown in **(F)** and were used for visualization only, not for quantification. Scale bar = 1 cm. **(F)** Quantification of bacterial titer in WT, *fec*, *anac017-1*, and *rao2-1* 72 h after infiltration. n = 16 biological replicates; error bars represent minimum to maximum values. Statistical differences for all panels were tested by one-way ANOVA followed by Tukey’s multiple comparisons test (P < 0.05).

### PTI signaling triggers an RBOH-independent, intracellular H_2_O_2_ signature

Mitochondria-derived ROS were recently pinpointed as the primary trigger of ANAC017-mediated retrograde signaling (*21*) (Fig. 3A). We hence assessed PTI-associated intracellular ROS and redox dynamics with the genetically encoded H_2_O_2_ biosensors HyPer7 and roGFP2-Orp1 in the cytosol and the nucleus of Arabidopsis leaves, using plate reader fluorimetry and confocal microscopy (*3*, *22*, *23*) (Fig. 3B). Both sensors showed pronounced, sustained oxidation in response to 1 µM flg22, indicating major intracellular H_2_O_2_ flux. Their signatures were similar overall, typically with two oxidation phases before recovery to baseline after at least 8 hours (Fig. 3C, D; Fig. S2A-L). Oxidation was apparent already in the first measurement interval, i.e. within 2 min, indicating fast intracellular H_2_O_2_ release. The glutathione redox potential (*E*_GSH_) biosensor Grx1-roGFP2 showed a similar signature (Fig. 3E; Fig. S2M-R), consistent with glutathione-based H_2_O_2_ detoxification in the cytosol (*24*). Confocal imaging revealed that oxidation signatures occur across cells and tissue types of the leaf (Fig. 3F-H; Movie S1, S2). The signatures followed a dose-response relationship across four orders of magnitude and remained detectable at 1 nM flg22 (Fig. S2). While the signatures were present in all experiments and their shapes were comparable within the individual experiment enabling reliable comparison, they showed a degree of variability in their specific shapes between experiments (Fig. S3).

**Figure 3.**
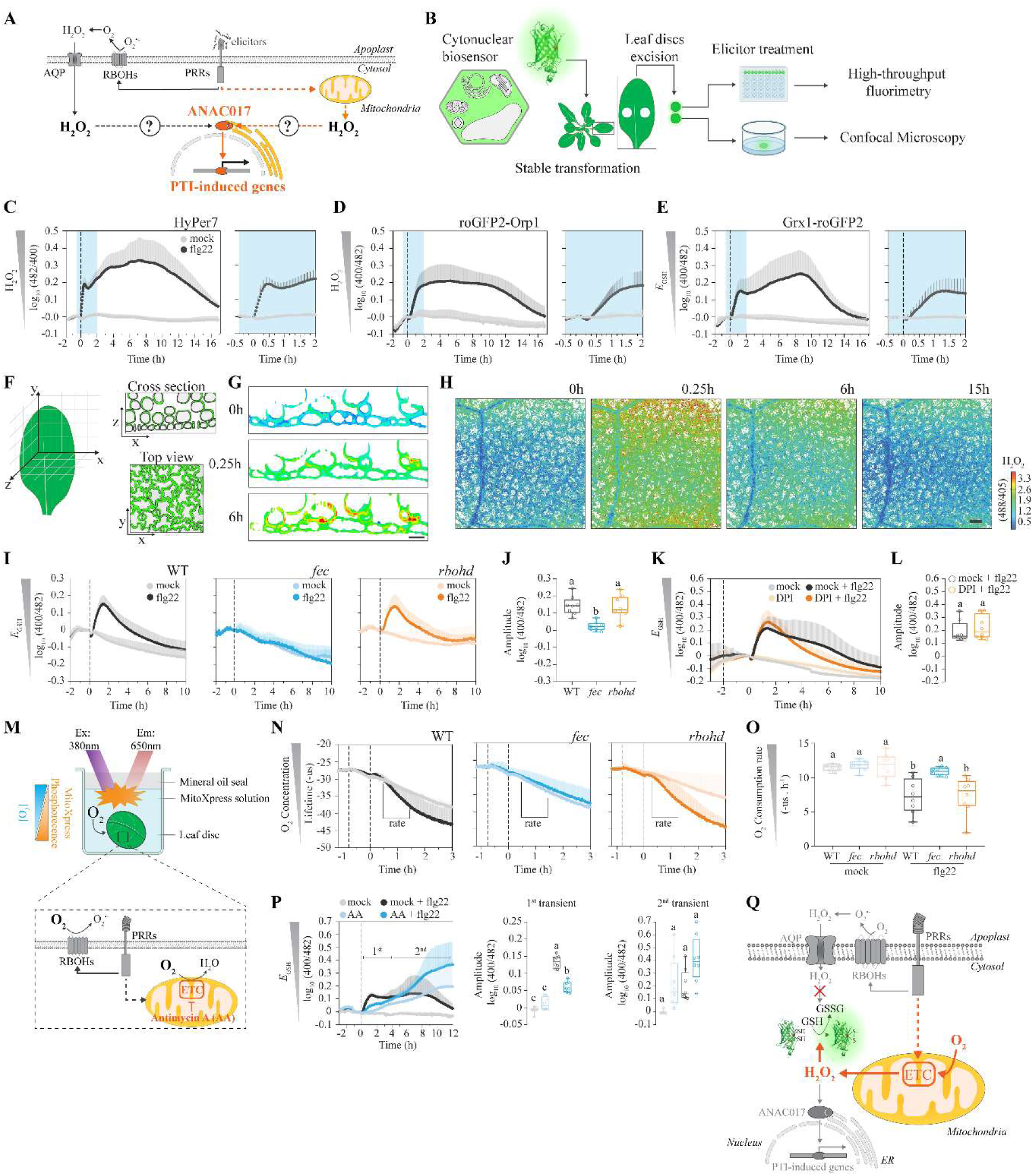
Mitochondrial-derived H₂O₂ contributes to PTI signaling. **(A)** Schematic representation of the hypothesized sources and signaling functions of H₂O₂ during PTI. **(B)** Experimental approach for monitoring cytosolic and nuclear physiological parameters during PTI using genetically encoded biosensors. Biosensor lines were generated by stable transformation and analyzed in leaf discs following elicitor treatment using high-throughput fluorimetry or confocal microscopy. **(C–E)** Time course of redox dynamics following treatment with 100 nM flg22 or water (mock). H₂O₂ dynamics were monitored in the cytosol using **(C)** HyPer7 and the cyto-nuclear compartment using **(D)** roGFP2-Orp1, while glutathione redox potential *E*_GSH_ was monitored in the cyto-nuclear compartment using **(E)** Grx1-roGFP2. Dashed lines indicate the time of flg22 or mock application. The blue-shaded regions in the left panels correspond to the magnified time intervals shown in the right panels. n = 12 biological replicates; lines represent mean values; error bars represent SD. **(F)** Schematic representation of the spatial sampling strategy for confocal imaging of leaf discs, including cross-sectional and top-view imaging. **(G)** Representative confocal images showing H₂O₂ dynamics in the leaf epidermis following 1 µM flg22 treatment at 0, 0.25, and 6 h monitored using HyPer7. Scale bars = 100 µm. **(H)** Representative spatial maps of H₂O₂ dynamics following 1 µM flg22 treatment at 0, 0.25, 6, and 15 h. H₂O₂ levels were monitored using HyPer7. Scale bar = 50 µm. **(I)** Time course of *E*_GSH_ dynamics in WT, *fec*, and *rbohd* following treatment with 100 nM flg22 or water (mock), monitored using Grx1-roGFP2. Dashed lines indicate the time of flg22 application. Lines represent mean values; n = 8 biological replicates; error bars represent SD. **(J)** Quantification of the amplitude of the flg22-induced *E*_GSH_ response in WT, *fec*, and *rbohd*. Lines represent mean values; n = 8 biological replicates; error bars represent minimum to maximum values. **(K)** Effect of DPI treatment on the flg22-induced *E*_GSH_ response monitored using Grx1-roGFP2. Plants were treated with 1 µM DPI or 0.1% DMSO (mock) (first dashed line) prior to treatment with 100 nM flg22 or water (mock) (dashed line at x = 0 h). Lines represent mean values; n = 8 biological replicates; error bars represent SD. **(L)** Quantification of the amplitude of the *E*_GSH_ response following DPI and flg22 treatment. n = 8 biological replicates; error bars represent minimum to maximum values. **(M)** Schematic representation of the experimental setup used to monitor oxygen consumption in leaf discs using the MitoXpress Xtra oxygen-sensitive probe. **(N)** Oxygen consumption dynamics in WT, *fec*, and *rbohd* following treatment with 100 nM flg22 or water (mock). Oxygen consumption was determined from the rate of change in MitoXpress fluorescence lifetime. First dashed line indicates well sealing with mineral oil while second dashed line (x = 0 h) indicates 100 nM flg22 or water (mock) treatment. Insets indicate the regions used to calculate oxygen consumption rates. Lines represent mean values. n = 8 biological replicates; error bars represent SD. **(O)** Quantification of oxygen consumption rates in WT, *fec*, and *rbohd* following mock or flg22 treatment. n = 8 biological replicates; error bars represent minimum to maximum values. **(P)** Effect of antimycin A (AA) on the flg22-induced *E*_GSH_ response. Time courses of *E*_GSH_ following mock or flg22 treatment in the presence or absence of AA are shown on the left. First dashed lines indicate 50 µM AA or mock (0.1% ethanol) treatment while second dashed lines (x = 0 h) indicate 100 nM flg22 or mock (water) treatment. Lines represent mean values; n = 8 biological replicates; error bars represent SD. Quantification of the amplitudes of the first and second transient responses is shown on the right. Error bars represent minimum to maximum values. **(Q)** Proposed model for the contribution of mitochondrial-derived H₂O₂ to PTI signaling. Elicitor perception activates RBOHD-dependent apoplastic H₂O₂ production and a parallel intracellular pathway leading to mitochondrial H₂O₂ production. Mitochondrial-derived H₂O₂ contributes to the cytosolic glutathione redox response and activates ANAC017-dependent expression of PTI-induced genes. Statistical differences for all panels were tested by one-way ANOVA followed by Tukey’s multiple comparisons test (P < 0.05).

The intracellular oxidation signature required FLS2-signaling since it was absent in *fec* (Fig. 3I). We hypothesized that its first phase might reflect RBOHD-activation, since the extracellular ROS burst occurs in a similar timeframe (Fig. 2D; Fig. S4A) and ROS are believed to enter the cytosol. Yet, the signatures were maintained in an *rbohd* knockout line (Fig. 3I, J; Fig. S4A, B) and in the presence of the RBOH-inhibitor diphenyleneiodonium chloride (DPI; Fig. 3K, L; Fig. S4A, B). Those data reveal that pronounced intracellular ROS and redox signatures are triggered as part of PTI, yet independently of the RBOH-dependent extracellular ROS burst. This conclusion is in line with recent observations of similar H_2_O_2_ dynamics in response to flg22 and elf18 (*4*), which were found to be dependent on the co-receptor BRI1-associated kinase (BAK1) and the receptor-like cytoplasmic kinase BOTRYTIS-INDUCED KINASE 1 (BIK1), but not RBOHD, as with an RBOH-independent ROS burst during *P. parasitica* infection (*15*).

### The H_2_O_2_ signature requires mitochondrial respiration

Since mitochondria-derived ROS activate ANAC017 (*21*), H_2_O_2_ production by the respiratory chain was a candidate source of PTI-associated intracellular ROS signature. The oxygen consumption rate of Arabidopsis leaves increased sharply upon flg22 exposure in an FLS2-dependent manner, being absent in *fec* and *bak1-5 bkk1-1* backgrounds (Fig. 3M-O, Fig. S4C, D). The increase persisted in *rbohd* (Fig. 3N, Fig. S4A, B) suggesting that RBOHD-mediated superoxide production contributes to a minor extent to the burst in oxygen consumption. Of note, those data highlight that historical naming of the NADPH oxidase enzymes in plants as ‘Respiratory burst oxidase homologues’, i.e. RBOH, is potentially misleading. Instead, the data pinpoint mitochondrial respiration, the dominant cellular oxygen sink, as the likely mechanism. Because respiratory complexes are major sites of ROS production (*25*, *26*), we impaired respiration with electron transport inhibitors and decreased O_2_ availability. The complex III inhibitors antimycin A and myxothiazol suppressed the flg22-induced Grx1-roGFP2 signature in leaf tissue in the dark (Fig. 3P; Fig. S5A, B). While antimycin A alone caused a steady rate of oxidation, the first phase of the flg22 response was inhibited. The complex IV inhibitor cyanide and the alternative oxidase (AOX) inhibitor *n*-propyl gallate (nPG) dampened the redox signature (Fig. S5E-J), as did lowering atmospheric oxygen levels (Fig. S5K, L). Genetic impairment of complex II activity (*dsr1* and *sdhaf2*) retained the initial oxidation but suppressed later phases, suggesting specific contributions of respiratory paths to the signature (Fig. S5M, N). Mitochondrial respiratory flux is thus required for the elicitor-induced redox signature (Fig. 3Q).

### An NAD redox signature is an ancient immune metabolic hallmark of PTI

We next sought the cause of increased mitochondrial ROS production. Because oxygen consumption increased (Fig. 3M-O, Fig. S4C, D), inhibited electron transport was unlikely. Alternatively, insufficient electron transport capacity may arise from increased reduced substrate and metabolic redox pressure (*25*). To assess the dynamics of the key ETC substrate NADH, we used the NAD redox state biosensor, Peredox-mCherry (*27*, *28*), expressed with cyto-nuclear localization, where it responds sensitively to changes in mitochondrial electron transport (*28–30*) (Fig. 4A). Flg22 triggered a fast NAD reduction peaking after about 1 h (Fig. 4B), followed by a partial re-oxidation, a milder second reduction phase and recovery after 8-10 h (Fig. S6A, B). The amplitude of the initial transient showed a strict dose-response relationship with flg22 concentration (Fig. S6A, B), similar to the H_2_O_2_ and *E*_GSH_ sensor responses. The NAD redox signature was extremely robust and thus represents a hitherto not recognized metabolic hallmark of PTI. Like the H_2_O_2_, it occurred across the entire leaf sample and was not specific to individual cell or tissue types (Fig. 4C, D). It was not specific to Arabidopsis, since a similar transient was triggered by flg22 also in tobacco (*Nicotiana tabacum*; (*31*)) leaf tissue and even moss (*Physcomitrium patens*) protonema after chitosan treatment (Fig. 4E). Its presence in a non-vascular plant suggests that the NAD redox transient represents an ancient PTI hallmark that evolved early in land plant evolution if not before. Similar to the H_2_O_2_ and *E*_GSH_ responses, the NAD transient was maintained in *rbohd* and with DPI, suggesting that it was not associated with NADPH oxidase activity (Fig. 4F, G).

**Figure 4.**
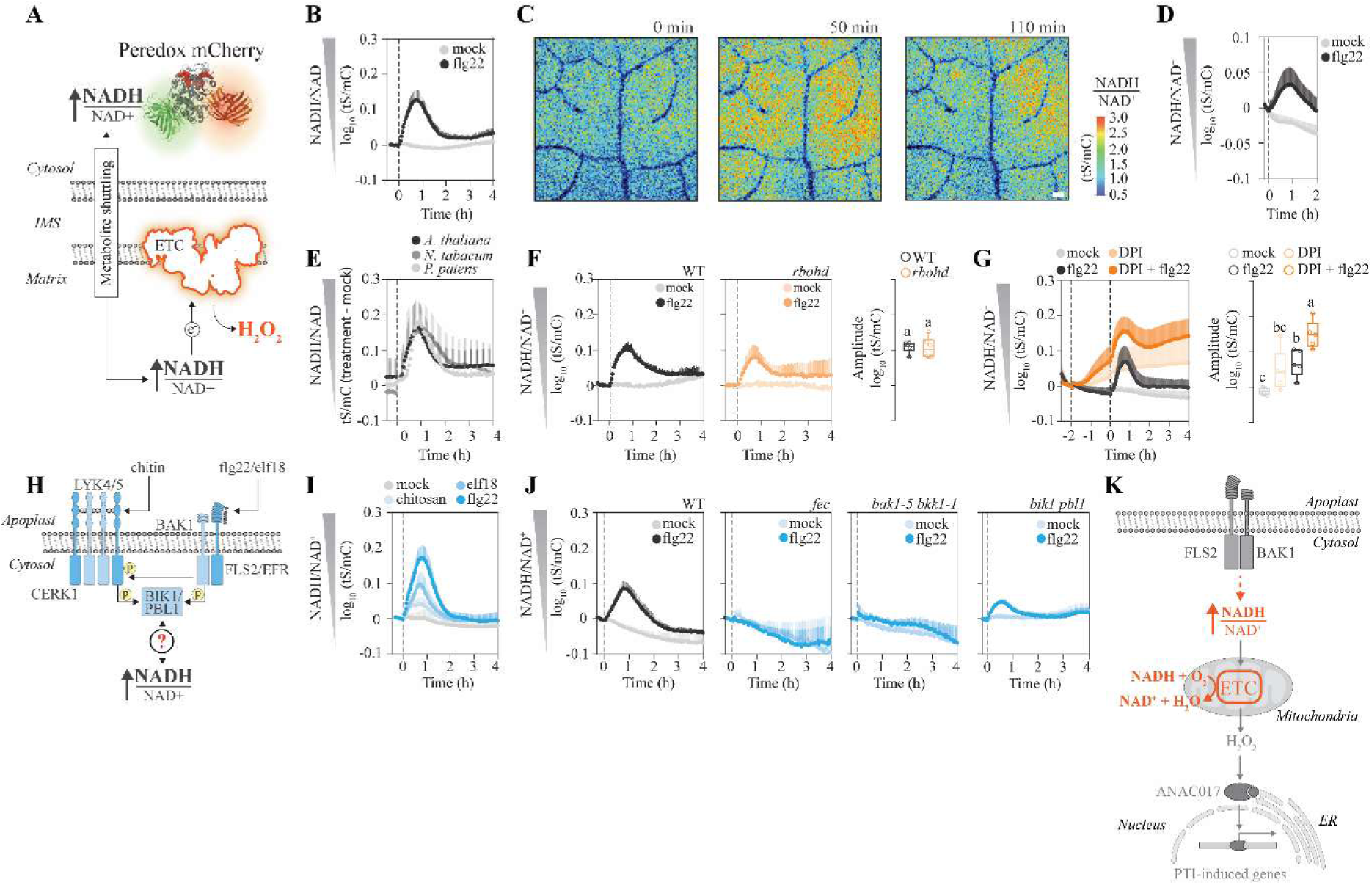
PTI induces a rapid reduction of the cytosolic NAD redox state. **(A)** Schematic representation of the hypothesized link between the PTI-induced H_2_O_2_ response and NAD redox state. Elicitor perception would increase the supply of reducing equivalents (increase NADH/NAD^+^ ratio) to the mitochondrial electron transport chain resulting in increased mitochondrial H₂O₂ production. **(B)** Time course of cyto-nuclear NAD redox dynamics in Arabidopsis leaf discs expressing the Peredox-mCherry biosensor following treatment with 100 nM flg22 or water (mock). Lines represent mean values; n = 5 biological replicates; error bars represent SD. **(C)** Representative confocal images showing spatial dynamics of the cyto-nuclear NAD redox state following flg22 or water (mock) treatment at the indicated time points. **(D)** Quantification derived from **(C)**. Lines represent mean values; n = 3 biological replicates; error bars represent SD. **(E)** Cyto-nuclear NAD redox dynamics following elicitor treatment in *Arabidopsis thaliana* (flg22 100 nM), *Nicotiana tabacum* (100 nM flg22), and *Physcomitrium patens* (0.1 mg / mL chitosan). Lines represent mean values; n = 12 biological replicates; error bars represent SD. **(F)** Cytosolic NAD redox dynamics following flg22 treatment in WT and *rbohd* plants (left). Quantification of the amplitudes of the first NAD reduction phase in response to flg22. The first reduction phase was defined as the maximum increase in the NADH/NAD⁺ ratio between 0 and 2 h after treatment. Dashed lines for all previous panels indicate elicitor treatment of either 100nM flg22 or 0.1 mg / mL chitosan. **(G)** Time course of cytosolic NAD redox dynamics in the presence or absence of DPI (left). First dashed line indicates treatment with 1 µM DPI or 0.1% DMSO (mock) while second dashed line (x = 0 h) indicates treatment with 100 nM flg22 or water (mock). Quantification of the amplitudes of the first NAD reduction phase in response to flg22 or mock in the presence or absence of DPI (right). The first reduction phase was defined as the maximum increase in the NADH/NAD⁺ ratio between 0 and 2 h after treatment. n = 5 biological replicates; error bars represent minimum to maximum values. **(H)** Schematic representation of the proposed connection between pattern-recognition receptor signaling and the PTI-induced cytosolic NAD redox response. Perception of elicitors by distinct receptors activates downstream signaling components, leading to a rapid reduction in the cytosolic NAD redox state. LYK4/5, LYSM-CONTAINING RECEPTOR KINASE 4 and 5; CERK1, CHITIN ELICITOR RECEPTOR KINASE 1; BAK1, BRI1-ASSOCIATED KINASE 1; FLS2, FLAGELLIN SENSING 2; EFR: LRR-RK EF-TU RECEPTOR; BIK1, BOTRYTIS-INDUCED KINASE 1; PBL1, serine/threonine-protein kinase PBL1; elf18, 18-amino-acid epitope derived from bacterial elongation factor Tu; flg22, 22-amino-acid epitope derived from bacterial flagellin. **(I)** Cyto-nuclear NAD redox dynamics in WT plants following treatment with the indicated elicitors. Dashed line indicates elicitor treatment. Chitosan = 0.1 mg / mL, elf18 and flg22 = 100 nM. **(J)** Cytosolic NAD redox dynamics in WT, *fec*, *bak1-5 bkk1-1* and *bik1 pbl1* following treatment with 100nM flg22. n = 8 biological replicates. Lines represent mean values. Error bars represent SD. **(K)** Proposed model linking elicitor perception, metabolic activation, mitochondrial respiration, and the cytosolic NAD redox response during PTI. Elicitor perception activates central carbon metabolism and increases the supply of reducing equivalents to the mitochondrial electron transport chain, resulting in a rapid reduction of the cytosolic NAD redox state and increased mitochondrial H₂O₂ production. Mitochondrial H₂O₂ subsequently contributes to ANAC017-dependent induction of PTI-associated genes. Statistical differences for all panels were tested by one-way ANOVA followed by Tukey’s multiple comparisons test (P < 0.05).

### The NAD redox transient is triggered downstream of the receptor-like kinase module and upstream of mitochondrial respiration

The NAD redox transient was activated by different elicitors (flg22, elf18 and chitosan) sensed by different PRRs, suggesting activation downstream of the specific receptors (Fig. 4H, I). Receptor-like kinase (RLK) signaling was nevertheless essential, since the transient was absent in *fec* and *bak1-5 bkk1-1* backgrounds upon flg22 treatment (Fig. 4J). It was only partially inhibited in *bik1 pbl1* (Fig. 4J), suggesting that RLK-mediated PTI-signaling activates central redox metabolism beyond RBOH activation, with a signal upstream of BIK1/PBL1 likely contributing. BAK1-mediated activation of the sugar transporter STP13 by phosphorylation has provided a blueprint for how RLK-based PTI may connect with metabolic regulation (*5*, *6*) and flg22 induces phosphorylation of several players in central carbon metabolism (*32*, *33*). To test the role of the mitochondria in shaping the NAD redox transient, we could not titrate electron transport with inhibitors without severely reducing the NAD pool and saturating the NAD redox sensor (Fig. S6C), and therefore titrated oxygen availability instead. Atmospheric oxygen levels of 10 and 5% did not affect baseline NAD redox status but enhanced the flg22-induced transient (Fig. S6D-F), validating a role for respiration in shaping it through NADH oxidation (Fig. 4K). How the reduction of the NAD pool is initiated, however, remained unexplained.

### Activation of respiratory carbon metabolism and sugar mobilization is required for PTI

To understand the cause of the NAD redox transient, we focused on high-capacity oxidative routes capable of sudden reductant mobilization. Prior work had suggested a BAK1-dependent increase of glucose availability in the cytosol upon PTI (*5*, *6*), but potential metabolic consequences had remained undefined. External sucrose, fructose, or glucose reduced the cyto-nuclear NAD pool, possibly due to sugar uptake into the cytosol and breakdown via glycolysis and the TCA cycle (Fig. 5A, B). In leaf tissue expressing the cytosolic glucose sensor FLIP Gluc 600 µM (*34*), flg22 triggered a rapid transient of cytosolic glucose depletion that largely mirrored the initial NAD redox transient (Fig. 5C). To confirm that depletion reflected increased catabolic flux rather than decreased mobilization, we used ^13^C-glucose feeding (Fig. 5D). ^13^C incorporation into sucrose increased under control conditions but was diminished by flg22 (Fig. 5E, F), showing that labeled glucose was efficiently metabolized by other pathways upon elicitation that outcompeted sucrose synthesis. Metabolite profiling at the first NAD reduction peak (50 min) and after partial recovery (120 min) confirmed glucose and sucrose depletion in response to flg22 (Fig. 5G, H; Fig. S7; Table S4). Increased pyruvate and alanine indicated enhanced glycolytic activity. GABA and succinate were also accumulated, reflecting increased NADH/NAD^+^, while the OAA-derived amino acids Asp and Lys were decreased. None of those changes occurred in *fec*, demonstrating PRR-signaling as the cause (Fig. 5G). Inhibiting hexokinase, the first glycolytic enzyme, with 2-deoxyglucose (2-DG), decreased the NAD redox signature in a dose-dependent manner down to complete suppression (Fig. 5I), and mannose had the same effect (Fig. S8A, B), demonstrating that glycolytic flux is essential. Hexokinase inhibition also abolished the flg22-induced burst in oxygen uptake (Fig. 5J), the *E*_GSH_ signature (Fig. 5K; Fig. S8C, D) and the response of MDR- and PTI-associated transcripts (Fig. 5L, M). Sugar catabolism via glycolysis is therefore strictly required for the intracellular respiratory burst. 2-DG also slowed down the extracellular ROS burst without changing total ROS released (Fig. S9), suggesting that sugar catabolism influences RBOH activity, while RBOH activity had no impact on the intracellular respiratory burst.

**Figure 5.**
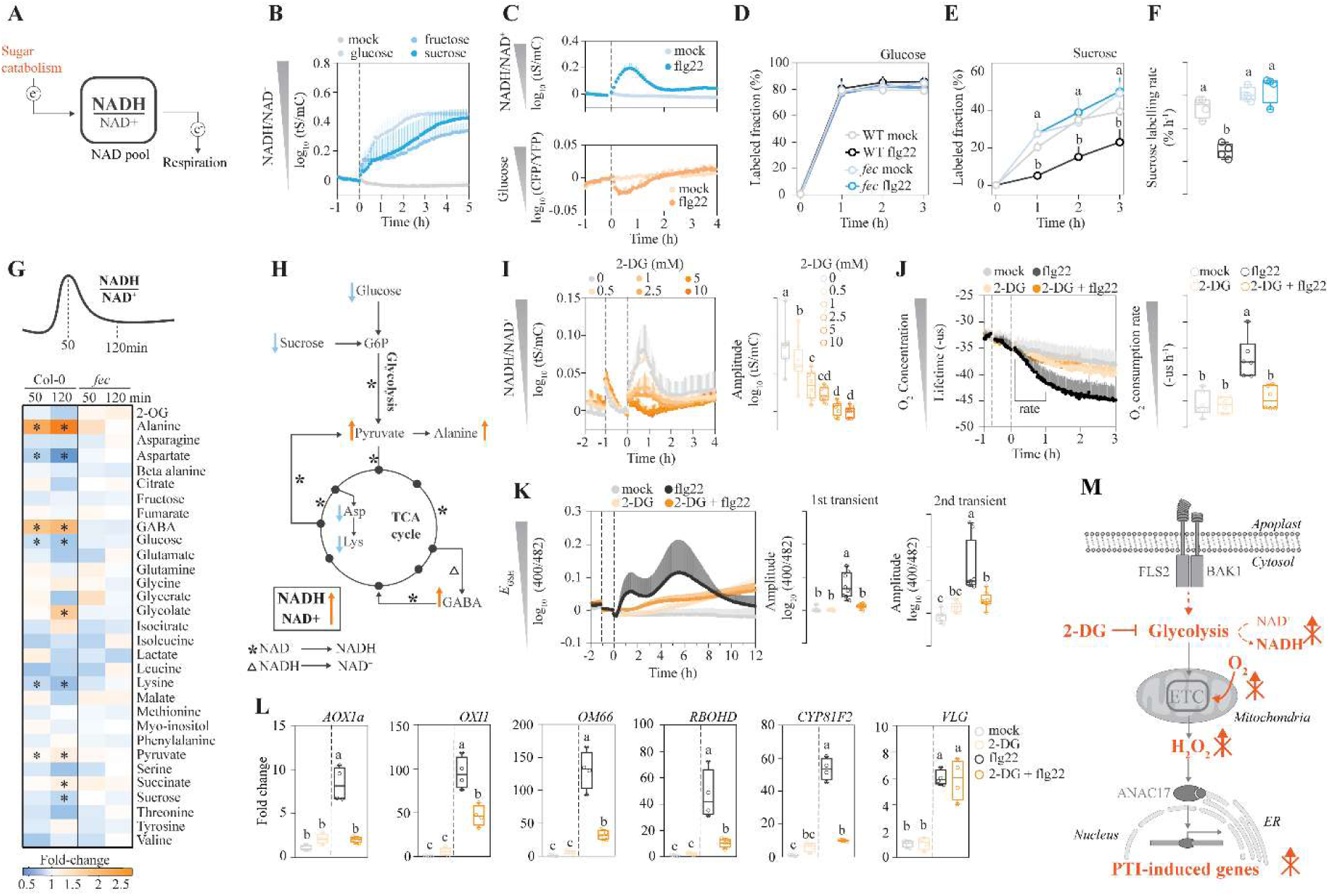
Pharmacological disruption of glycolysis attenuates the intracellular respiratory burst during PTI. **(A)** The cytosolic NAD redox state reflects the balance between electron supply from central carbon metabolism and electron efflux through respiration. **(B)** Time course of cytosolic NAD redox dynamics in Arabidopsis leaf discs expressing the Peredox-mCherry biosensor following treatment with 10 mM glucose, sucrose or fructose compared to water (mock). n = 6 biological replicates. Lines represent mean values. Error bars represent SD. **(C)** Parallelized time courses of cytosolic NAD redox (top) and glucose concentration (bottom) following addition of 100 nM flg22 in Arabidopsis thaliana leaf discs expressing either Peredox-mCherry or FLIP Gluc 600 µM. n = 8 biological replicates. Lines represent mean values. Error bars represent SD. GC-MS quantification of the fraction of isotope labelled **(D)** glucose or **(E)** sucrose 0, 1, 2 and 3 h after treatment with 100 nM flg22 in WT and *fec* mutant. **(F)** Labelling rate of sucrose after treatment with 100 nM flg22 in WT and *fec.* (D-F) n = 4 biological replicates. Error bars represent SD. Leaf discs were labelled with 5 mM 13^C^-glucose as described in the methods section. **(D-F)** Statistical differences were tested by two-way ANOVA followed by Tukey’s multiple comparisons test (P < 0.05). **(G)** GC-MS untargeted metabolomics of WT and *fec* after 50 and 120 min treatment with 1 µM flg22. Values shown are the fold-change between 1 µM flg22 and water (mock) treated samples. n = 8 biological replicates. Asterisks indicate statistical significance calculated using two-way ANOVA followed by Tukey’s multiple-comparisons test. p-values were adjusted for multiple testing using the Benjamini–Hochberg false discovery rate (FDR) correction. An adjusted p-value (q-value) < 0.05 was considered statistically significant. **(H)** Schematic representation of the observed changes in metabolite abundance shown in **(G)**. **(I)** Time courses of cytosolic NAD redox dynamics in Arabidopsis thaliana leaf discs pre-treated with 0, 0.5, 1, 2.5, 5 and 10 mM 2-deoxyglucose (2-DG) followed by 100 nM flg22 treatment. First dashed line indicates 2-DG treatment and second dashed line (x = 0 h) indicates flg22 treatment. Lines represent mean values. Error bars represent SD. Quantification of the amplitudes of the first NAD reduction phase in response to flg22 in leaf discs pre-treated with 2-DG (right panel). The first reduction phase was defined as the maximum increase in the NADH/NAD⁺ ratio between 0 and 2 h after treatment. Error bars represent minimum to maximal values. n = 8 biological replicates. **(J)** Oxygen consumption dynamics in WT following pre-treatment with 10 mM 2-DG or water (mock) and subsequent treatment with 100 nM flg22 or water (mock). First dashed line indicates well sealing with mineral oil while second dashed line (x = 0 h) indicates 100 nM flg22 treatment. 2-DG treatment was done at the beginning of the time-course. Lines represent mean values. Insets indicate the regions used to calculate oxygen consumption rates. Oxygen consumption was determined from the rate of change in MitoXpress fluorescence lifetime. The right panel shows quantification of oxygen consumption rates in WT after the abovementioned treatments. n = 6 biological replicates. Error bars represent minimum to maximum values. **(K)** Time courses of cytosolic *E_GSH_* dynamics in Arabidopsis leaf discs pre-treated with 10 mM 2-deoxyglucose (2-DG) or water (mock) followed by 100 nM flg22 or water (mock) treatment. First dashed line indicates 2-DG treatment and second dashed line (x = 0 h) indicates flg22 treatment. Lines represent mean values. The right panel shows the quantification of the first and second *E_GSH_* transient amplitude calculated from the left panel. The first transient was defined as the maximum response between 0–2 h and the second transient as the maximum response between 2–12 h. Error bars represent minimum to maximum values. n = 8 biological replicates. **(L)** Relative transcript abundance of *AOX1a*, *OXI1*, *OM66*, *RBOHD*, *CYP81F2*, and *VLG* in leaves of 4-week-old Arabidopsis thaliana plants. Leaf discs were pre-treated with 10 mM 2-DG or mock for 1 h before elicitation. Samples were collected 50 min after treatment with 100 nM flg22 or water (mock). Transcript levels were normalized to WT mock control. n = 4 biological replicates; error bars represent minimum to maximum values. **(M)** Proposed model for the role of glycolytic activation in the intracellular respiratory burst during PTI. Elicitor perception stimulates sugar catabolism through glycolysis, increasing the supply of reducing equivalents to the mitochondrial electron transport chain and promoting mitochondrial oxygen consumption and H₂O₂ production. Mitochondria-derived H₂O₂ contributes to the cytosolic glutathione redox response and ANAC017-dependent induction of PTI-associated genes. Statistical differences for all panels except **(D-G)** were tested by one-way ANOVA followed by Tukey’s multiple comparisons test (P < 0.05).

### The redox metabolic branch of PTI signaling integrates carbon metabolic status

We next challenged the intracellular respiratory burst model with genetic interventions in carbon metabolism. To perturb the flexibility of mitochondrial carbon metabolism, we used a mutant lacking NAD-malic enzyme activity (NAD-ME1 and NAD-ME2) (*35*, *36*). The nad-*me1-2 2-3* background showed a lower NAD reduction amplitude (Fig. S10A) and the *E*_GSH_ signature was suppressed (Fig. S10B), while the extracellular ROS burst was unchanged (Fig. S10C). By contrast, mutants lacking key mitochondrial enzymes in NAD concentration homeostasis (NADKc) and uncoupling (UCP1, AOX1a) did not show changes in their signatures (Fig. S10D, E). We further assessed the impact of modified intracellular free sugar availability using the *stp1 stp13* mutant line, which was recently found to have impaired flg22-induced uptake of apoplastic hexoses (*5*, *6*). This mutant showed no detectable changes in its NAD redox or *E*_GSH_ signatures (Fig. S11A-C), suggesting that this route of cytosolic sugar uptake does not make any major quantitative contribution to oxidative metabolism and signaling. In contrast, *sweet11 sweet12*, which accumulates high levels of intracellular soluble sugars (*37*) showed enhanced amplitudes of the NAD redox signature and of the *E*_GSH_ signature in its first phase (Fig. S11D-F).

### Daytime-specific immunity is modulated by the redox metabolic branch of PTI signaling

The implication of these observations, i.e. that the PTI-associated signatures are shaped by carbon metabolic flux status, prompted us to assess the impact of the normal day-night cycle, over which leaf carbon status varies drastically (*38*). The NAD redox signature was similar at mid and end of day, but decreased markedly towards dawn, when carbohydrate resources are depleted, and was further diminished by starvation during an extended night (Fig. 6A, B). This striking correlation and the observation of decreased bacterial titers in the absence of ANAC017-mediated mitochondrial retrograde signaling (Fig. 2E, F) prompted us to carry out daytime-specific quantitative infection assays with bioluminescent *Pseudomonas syringae* pv. tomato DC3000 (*20*). Arabidopsis leaves showed the strongest bacterial titers after infection at midday. Infection success decreased gradually thereafter and reached a minimum at dawn (Fig. 6C, D); extended night did not diminish bacterial titers any further. Daytime-specific differences were absent in *fec*, showing that they depend on PTI signaling, rather than on direct effects of carbohydrate status on the bacteria (Fig. 6C, D). The intracellular respiratory burst integrates context-dependent metabolic status in Arabidopsis. Whether this role was adopted in early land plant evolution or represents a derived function deserves future investigation.

**Figure 6.**
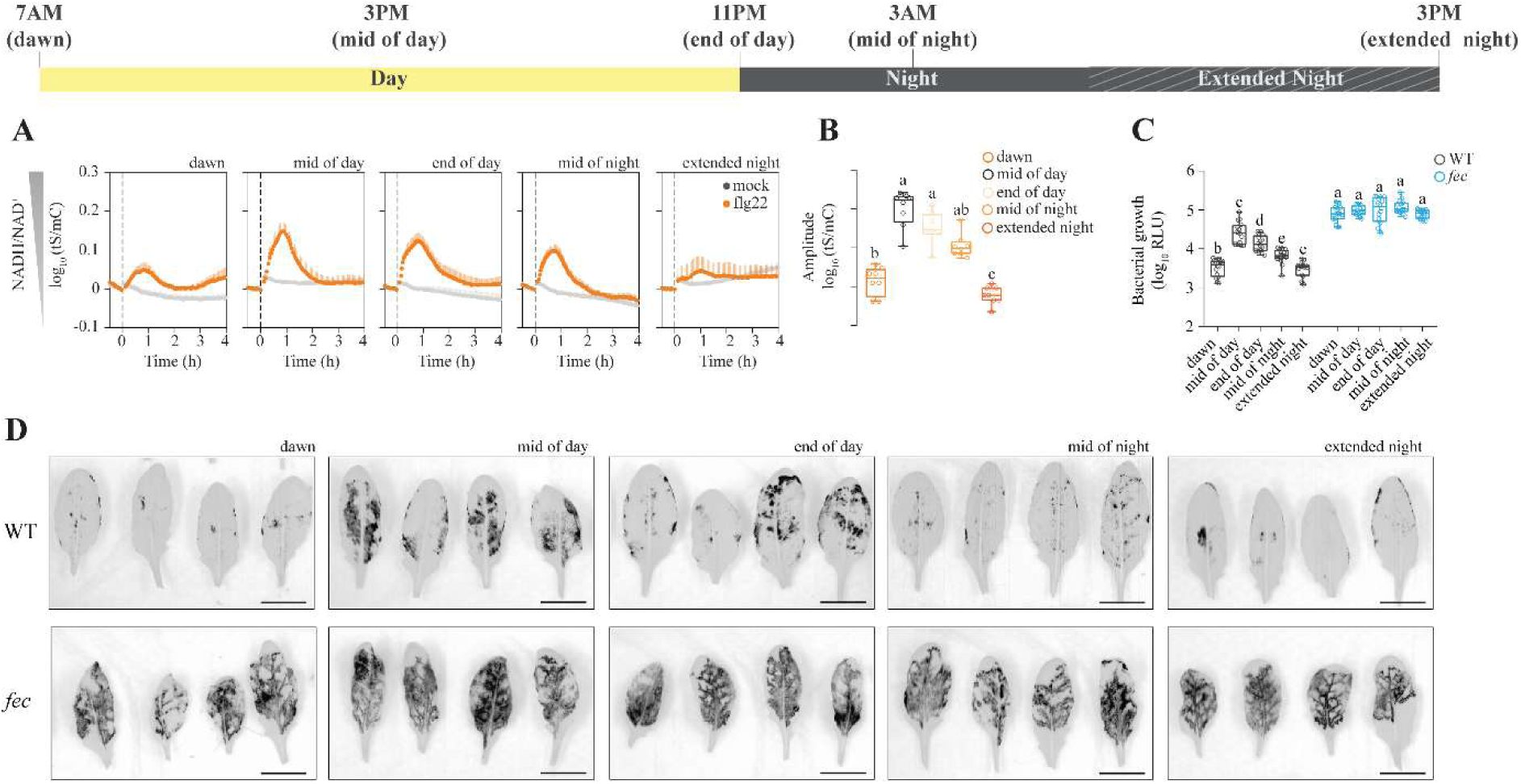
Susceptibility to bacterial pathogens and induction of the intracellular respiratory burst during PTI is daytime dependent. **(A)** Time course of cytosolic NAD redox dynamics in Arabidopsis leaf discs expressing the Peredox-mCherry biosensor following treatment with 100 nM flg22 or water (mock) at different times of the day. Leaf discs were excised at the times represented in the above scale bar. Treatments were carried out after 2 h 30 min dark incubation. Dashed lines indicate flg22 treatment. **(B)** Quantification of the NAD redox transient derived from (A). n = 8 biological replicates. Error bars represent minimum to maximum values. The first reduction phase was defined as the maximum increase in the NADH/NAD⁺ ratio between 0 and 2 h after treatment. Statistical differences were tested by one-way ANOVA followed by Tukey’s multiple comparisons test (P < 0.05). **(C)** Quantification of bacterial growth in WT and *fec* 72 h after infiltration with *Pseudomonas syringae* pv. tomato DC3000 (OD = 0.01) carrying a luciferase operon at different times of the day. n = 12 biological replicates. Times of infiltration are represented in the above scale bar. Statistical differences were tested by two-way ANOVA followed by Tukey’s multiple comparisons test (P < 0.05). **(D)** Representative chemiluminescent images of (C). Images are representative of the experiment shown in (C) and were used for visualization only, not for quantification. Scale bar = 1 cm.

## Discussion

Here, we describe a metabolic branch of PTI characterized by an intracellular respiratory burst (Fig. 1A). Genetically encoded biosensors resolved fast redox-metabolic changes in living leaf tissue and pinpoint mitochondria as central connectors of metabolic flux and defense signaling (*39*). Following the biosensing-based observation of an elicitor-induced intracellular H_2_O_2_ signature (*3*), Arnaud et al. (2023) (*4*) established that elicitors and *Pseudomonas syringae* result in biphasic cyto-nuclear H_2_O_2_ dynamics independently of apoplastic ROS production. We provide a unified framework for these signatures, a signaling and metabolic nexus bridging the plasma membrane, cytosol, mitochondria, ER, and nucleus. While we utilize an elicitor treatment model to reliably resolve redox metabolic signatures, those events are likely highly organized at the subcellular level during genuine plant-microbe encounters (*40*), for instance through immobilization and clustering of mitochondrial subpopulations and the ER at pathogen entry sites, and through the selective ROS release by those subpopulations (*13*). Mitochondrial ROS released in the direct vicinity of the ER membrane promote mitochondria-ER contacts, enabling ROS signal transfer to initiate the unfolded protein response (UPR) and bolster host immunity (*15*). This is a plausible framework for how mitochondria activate ANAC017-mediated retrograde signaling, with our findings identifying acute primary metabolic activation as the trigger of these redox changes. Mitochondrial retrograde signaling is a conserved stress module (*12*), and its early-identified association with defense (*11*) now gains a mechanistic basis. The intracellular respiratory burst may thus integrate several previous, mechanistically disconnected findings on mitochondria in host defense (*9*, *10*, *13*, *15*, *41–43*).

The independence of the intracellular respiratory burst from NADPH oxidases (Fig. 3I-L, N, O) aligns with recent observations (*4*, *15*). While apoplastic H₂O₂ is largely restricted to plasma membrane nanodomains by cytosolic scavenging systems (*44*), and chloroplasts also generate immunogenic ROS (*45–47*), our work establishes a non-redundant requirement for mitochondria (Fig. 3P, Fig. S5). Such coordinated mitochondrial bursts, on top of documented mitochondrial ROS production in biotic stress (*7*, *10*, *48*), may prime effector-triggered immunity and systemic acquired resistance. Although NAD homeostasis is a core determinant of resistance (*49*) and NAD-derived ‘info-metabolites’ are emerging signals (*50*, *51*), our study establishes a function for NAD as a metabolic redox carrier (Fig. 4). Future research will need to assess potential mechanistic intersections between both fundamental defense signaling strategies. Our model also redefines primary carbon metabolism as an active regulatory partner in defense (*52*). A recent study of viral infection showed parallels, including a shift in NAD redox status and the significance of mitochondrial NAD-malic enzyme (*53*). Furthermore, the rapid cytosolic glucose depletion we observe (Fig. 5C, G, H) occurs on a timescale that precedes the slower glucose uptake by STPs (*5*, *6*), suggesting that the activation of host-mediated sugar import primarily serves as an ancillary role, in agreement with our observation that *stp1 stp13* mutants did not affect the pathway significantly (Fig. S11A-C). Thus, while sugar status and mobilization have historically been implicated in defense through a passive source-to-sink carbon reallocation framework (*2*, *54*), we propose a model where acute glycolytic flux and mitochondrial retrograde signaling operate as a direct, instructive branch of plant pattern-triggered immunity (Fig. 5I-L).

Conservation of the intracellular respiratory burst between bryophytes and seed plants (Fig. 4E) suggests that cellular metabolic activation is an ancestral process in land plant immunity. Core PTI components, including the PBL/RBOH module and chitin receptor signaling, are conserved among land plants (*55*, *56*), whereas other immune pathways and specific elicitor receptors emerged later (*57*). Functional parallels extend to the respiratory metabolic reprogramming that drives immune cell activation in mammals, including macrophages, dendritic cells and T-cells (*58–61*).

Although the metabolic PTI branch negatively modulates host defense against the hemibiotrophic pathogen *Pseudomonas syringae* in our system, its function may differ depending on the patho-system, including (1) “starving” invaders by burning soluble carbohydrates, (2) mobilizing carbon resources for defense metabolite synthesis or (3) generating the ATP necessary to power acute cellular defense (even though we observed no obvious change in cyto-nuclear ATP concentration; Fig. S12). Those archetypical metabolic roles may have provided the basis to evolve a retrograde signal built upon this ancient metabolic reprogramming. Further, the metabolic PTI branch provides a physiological mechanism for daytime-dependent immunity, reinforcing recognized transcription- and redox-based circadian clocks that gate defense responses in the morning to minimize growth costs at night, when stomata are typically closed and invasion via the stomata is less likely to occur (*62–64*). Harnessing this mechanism of daytime-specific immunity may inform the development of advanced strategies for crop protection.

### Limitations and perspectives

How immune-activated kinase signaling affects enzymes of respiratory carbon metabolism remains to be resolved. Both, direct phosphorylation-based mechanisms (*65*) as well as indirect mechanisms via energy signaling offer testable hypotheses. Future investigations must also map which specific metabolic substrates are used and assess whether these conserved metabolic responses are similarly triggered during mutualistic microbe-host interactions to shape the plant microbiome.

## Acknowledgements

We thank Edward Smith, Thomas Nietzel, Janina Steinbeck for preliminary experimental exploration and Aurora Martin (Kaiserslautern) for the technical support. We are grateful to Jan-Riemer (Cologne), Bruce Morgan (Saarbrücken), Marc Knight (Durham, Halle), George Ratcliffe (Oxford), Andreas J Meyer (Bonn), Lee Sweetlove (Oxford), A. von Schaewen (Münster) and Malcolm Bennett (Nottingham) for critical discussion, and thank Beatriz Moreno-Garcia (Madrid) for the kind gift of the *N. tabacum* Peredox-mCherry line, Hirofumi Nakagami (Cologne) for the bioluminescent Pseudomonas strains and Wolf Frommer (Düsseldorf) for the glucose biosensor lines.

## Funding

This work was supported by the Deutsche Forschungsgemeinschaft (DFG) through the collaborative research grant PAK918 (V.G.M., M.Sch.; project no. 289357231), the DFG research unit RU5573 (M.Sch., project no. 507704013), the DFG research unit RU5640 (M.Sch., project no. 515385982), the DFG research grant (M.Sch., A.R.F. project no. 508398975), the Heisenberg Grant (T.I. project no. 463060822), the Walther-Benjamin postdoctoral grant (A.C.V. project number 560830917) and the infrastructure grants INST211/903-1 FUGG (M.Sch., project no. 429542710) and INST 211/1110-1 FUGG (T.I. project no. 529034606). P.B. was funded by National Council for Scientific and Technological Development (CNPQ) through a postdoctoral fellowship (project no. 200385/2022-4).

C.Z. is funded by the University of Zurich and the Gatsby Charitable Foundation.

## Author Contributions

Conceptualization: PB, EFA, LMV, MSch

Methodology: JON, SL, SW

Investigation: PB, EFA, LSK, ML, JON, MSz, GD, ACV, SR, CR, AS, JN, DD, KZ, AA, AC, RJ, MFS, JM

Visualization: PB, MSch

Funding acquisition: MSch

Project administration: MSch

Supervision: ANN, VGM, SMS, TI, ARF, LMV, MSch

Writing – original draft: PB, MSch

Writing – review & editing: PB, EFA, EL, ANN, VGM, SMS, TI, CZ, ARF, LMV, MSch

## Competing Interests

Authors declare that they have no competing interests.

## Data, Code, And Materials Availability

Untargeted metabolomics data obtained for this study are deposited at Metabolomics Workbench (ID: 8036). All additional data are available in the main text or the supplementary materials.

## Supplementary Materials

### Materials and Methods

#### Plant material and growth conditions

*Arabidopsis thaliana* Col-0 lines expressing fluorescent biosensors and knockout mutant lines used in this study are listed in Table S5. Knockout mutant lines used in this study were equipped with fluorescent protein biosensors using the floral dip method (*66*). At least two independent transgenic lines were selected based on antibiotic resistance and fluorescence for each mutant.

*Arabidopsis thaliana* and *Nicotiana tabacum* seeds were surface sterilized with 70% ethanol, plated in half-strength Murashige and Skoog medium (MS: Duchefa, Haarlem, the Netherlands) supplemented with 0.1% (w/v) sucrose, 0.05% (w/v) 2-(N-morpholino)ethanesulfonic acid (MES), adjusted to pH 5.8 (KOH), and 0.8% (w/v) plant agar (Duchefa, Haarlem, the Netherlands). Seeds were stratified in the dark at 4 °C for 48 h. Plant growth was performed in growth cabinets set to long day conditions (day: 16 h at 22 °C and 120 μE m^−2^ s^−1^, night: 8 h at 18°C). Seven days after transfer to light, seedlings were transferred to square plastic pots containing humidified soil (50% (v/v) VMV800 Vermehrungserde and 50% (v/v) EDE800 Einheitserde) (Balster Einheitserden, Frödenberg, Germany). Trays with pots were covered with a transparent plastic dome and placed back into the growth cabinet. After three days the dome was removed and plants were watered every two to three days with tap water. Plants were used for further experiments 4–5 weeks after germination (28–35 days old).

Protonema of *Physcomitrium patens* (Hedw.) Mitt. (*67*) (*Physcomitrium patens* (Hedw.) Bruch & Schimp., ecotype Gransden 2004; International Moss Stock Center (IMSC), accession number 40001) were grown axenically in liquid KNOP-ME medium as previously described (*68*). Cultures were regularly homogenized in fresh medium using an Ultra-Turrax T25 dispersion tool (IKA-Werke GmbH, Staufen, Germany) at 14,000 rpm for 1-1.5 min. Stable transgenic lines expressing Peredox-mCherry (*28*) under the control of the endogenous *Actin5* promoter and the *NOS* terminator (vector PTA2-Act5-Gateway) (*69*, *70*), integrated at the PTA2 locus via homologous recombination (*71*), were generated by polyethylene glycol-mediated protoplast transformation, as previously described (*72*). Cells were co-transformed with an uncut plasmid containing a *nptII* resistance cassette (pBsNNNEV) and selected for four weeks on geneticin G418 (12.5 µg/mL) before PCR-based genotyping for transgene presence and expression cassette integration at the PTA2 genomic locus (*68*).

### Transcriptomic meta-analysis

The 1,000 genes commonly upregulated by elicitor treatment (hereafter defined as Core-PTI transcripts) identified by Bjornson et al. (*16*) were used as the query set for Gene Ontology (GO) enrichment analysis. Complete GO annotations for the *Arabidopsis thaliana* genome were downloaded from the Gene Ontology database (downloaded on 06/12/2024). The annotation file was modified to include the 24 genes classified as the Mitochondrial Dysfunction Regulon (MDR) as a custom Biological Process term (*17*). Enrichment analysis was performed using g:Profiler (https://biit.cs.ut.ee/gprofiler/gost), with the Core-PTI transcripts as the query set and the modified GO annotation as the background. Enriched terms were considered significant at a Benjamini–Hochberg false discovery rate (FDR) < 0.05 (Table S1). MDR transcript responses were also evaluated individually for each elicitor (Table S2). The overlap between mitochondrial dysfunction and immune responses was further assessed using transcriptomic datasets from plants treated with the mitochondrial inhibitor antimycin A (*73*) or overexpressing ANAC017 (*19*) (Table S3). Venn diagrams were generated using the Bioinformatics and Systems Biology Tools web service (https://bioinformatics.psb.ugent.be/webtools/Venn/) and subsequently redrawn using Adobe Illustrator (Figure S1). To assess whether the observed overlaps exceeded those expected by chance, three random gene lists of equal size were sampled from the TAIR12 genome annotation, and the mean overlap was used as the expected overlap. Statistical significance was assessed using two-sided chi-square tests.

### RNA extraction and gene expression analysis

A total of 8 leaf discs were pooled from 4 different plants and transferred to 96-well plates containing 180uL of plate reader assay medium (10 mM MES, 10 mM MgCl_2_, 10 mM CaCl_2_, 5 mM KCl, pH 5.8 adjusted with KOH). Each pool of 8 leaf discs was considered one biological replicate. The plates were incubated in the dark for 2h 30 min before 20uL of either 1µM flg22 (100nM final concentration) or 20µL water (mock control) was added to each well. Leaf discs were collected 50 min after the treatment, dried using lint-free laboratory wipes (Kimwipes: Kimberly-Clark, Irving, TX, USA), and immediately flash-frozen in liquid nitrogen. Total RNA was isolated using the NucleoSpin RNA Plant kit, treated with DNase I (both from Macherey-Nagel, Düren, Germany). RNA (500 ng) was reverse transcribed using an iScript cDNA Synthesis Kit (BioRad, Hercules, CA, USA). SYBR Green detection (KAPA SYBR Fast qPCR Master Mix, Roche, Mannheim, Germany) in a C1000 Touch Thermal Cycler using the CFX 96 Real-Time System (Bio-Rad, Hercules, CA, USA) was used to perform RT–qPCR. The primers used for RT-qPCR are listed in Table S6. Gene expression was analyzed using the comparative Ct (ΔΔCt) method, with UBIQUITIN10 (UBQ10) used as the reference gene for normalization. Four biological replicates were analyzed per treatment/genotype, with three technical replicates per biological replicate.

### Apoplastic ROS burst assay

Measurement of the apoplastic ROS burst was performed as previously described (*3*). Leaf discs excised from 28d-old plants were submerged in water in 96-well plates overnight. The water was replaced with 0.5 mM L-012 (FUJIFILM Wako Chemicals U.S.A. Corp) in 10 mM MOPS/KOH (pH 7.4). Luminescence was measured in a CLARIOstar® microplate reader and after a short period of adaptation, 100 nM flg22 or water for control was added. The measurements were recorded for at least 1 hour after elicitor treatment. Where additional pre-treatments were performed prior to flg22 elicitation, these are indicated in the respective figure legends.

### Pseudomonas infection assays

Measurement of plant susceptibility to *Pseudomonas syringae* pv. *tomato* DC3000 carrying a luciferase operon (PtoLUX) was performed as previously described (*20*). A total of 10uL of a PtoLUX glycerol stock was transferred to 5mL of liquid King’s Broth medium containing rifampicin (50 µg / mL) and grown for 3 days at 28 °C with shaking at 150 rpm. After 3 days, 100µL of the culture was spread evenly in King’s Broth agar plates and incubated at 28 °C for additional 48 h. Subsequently, 2mL of 10mM MgCl_2_ was added to each plate to resuspend the PtoLUX cells, and the suspension was transferred to 2 mL tubes. The PtoLUX suspension was centrifuged at 4,000 rpm for 5 minutes, the supernatant was discarded, and the cells were resuspended in 2 mL of 10mM MgCl_2_. This washing step was repeated twice. The PtoLUX suspension was then diluted 1:10 in 10 mM MgCl₂, and the OD₆₀₀ was measured. The suspension was subsequently diluted to an OD₆₀₀ of 0.001 or 0.01. Approximately 1 h before infiltration, 1.5 L of water was added to the trays containing the plants to stimulate stomatal opening. PtoLUX suspensions at the indicated OD₆₀₀ or mock solution (10 mM MgCl₂) were infiltrated into four leaves per plant using a 1 mL syringe. The trays were covered with a plastic dome for 1 h after infiltration. After 48 or 72 h, leaf discs were excised from the infiltrated plants and transferred to white 96-well plates. Luminescence was measured in PtoLUX- and mock-infiltrated leaves using a CLARIOstar® plate reader with a 10 s integration time. Representative leaves were imaged using the chemiluminescence mode of an INTAS GelDoc system (INTAS Science Imaging Instruments GmbH, Göttingen, Germany). Chemiluminescence images were used for visualization only and not for quantification. Each biological replicate consisted of a pool of 11 leaf discs excised from four infected leaves of a single plant. A total of 12 or 16 biological replicates were used for each experiment, as indicated in the respective figure legends.

For PtoLUX infection assays performed at different times of day, starter PtoLUX cultures were prepared exactly 120 h before the respective infiltration time. For example, when infiltration was scheduled for 3:00 AM, the starter culture was prepared at 3:00 AM five days before infiltration. Accordingly, starter cultures were transferred to King’s Broth agar plates exactly 48 h before the scheduled infiltration time. Plants were grown under a 16 h light/8 h dark photoperiod, with the light period beginning at 7:00 AM and ending at 11:00 PM. Infiltrations were performed at 11:00 PM (end of day), 3:00 AM (mid of the night), 7:00 AM (end of night), 3:00 PM (mid of the day), or 3:00 PM following an extended dark period. For the extended-night condition, plant trays were covered with a black dome at 7:00 AM and kept in darkness until 3:00 PM, thereby extending the dark period by 8 h.

### Multi-well plate reader-based fluorimetry

Live monitoring of cytosolic physiology using genetically encoded biosensors was carried out in a CLARIOstar® microplate reader as previously described (*3*, *28*, *74*, *75*) with the following modifications. Each well of transparent 96-well plates was filled with 180 µL plate reader assay medium (10 mM MES, 10 mM MgCl_2_, 10 mM CaCl_2_, 5 mM KCl, pH 5.8 with KOH). Individual leaf discs excised from fully expanded leaves of 28-35-day-old plants were placed upside down into each well. A pair of soft tweezers was used to gently push the leaf disc towards the bottom of the well without damaging the central part of the disc. For *Physcomitrium patens*, protonemal cells were allowed to settle for 15 min in 50 mL tubes, after which the KNOP-ME medium was replaced with plate reader assay medium. A total of 200 µL of protonema cells was then transferred to 96-well transparent plates.

The 96-well plates containing the leaf discs were dark-adapted for 2.5 h inside the plate reader chamber pre-equilibrated at 25 °C, whereas plates containing *Physcomitrium patens* protonema cells were incubated overnight in the dark at 25 °C. Fluorescence intensity was measured in plate mode with top optics using a standard protocol with 35 excitation flashes per well and cycle, an orbital averaging with 3 mm diameter and 0.1 s settling time. Excitation and emission wavelengths were chosen for each biosensor: roGFP2-Orp1, Grx1-roGFP2 and HyPer7 : Ex = 400 ± 5 and 482 ± 8 nm/Em = 520 ± 20 nm; Peredox-mCherry: Ex = 400 ± 5 and 540 ± 10 nm/Em = 520 ± 20 and 610 ± 5 nm; FLIP Gluc 600µM and ATeam 1.03 nD/nA: Ex = 430 ± 10 nm/Em = 480 ± 10 and 530 ± 10 nm. Gain adjustments for each wavelength were done in the beginning of the study and kept the same for comparison. Fluorescence intensity was measured for about 2 h 30min before performing any treatment. Elicitor treatments and other chemical treatments were carried out by pipetting 20 µL of the corresponding solution into each well as described in figure legends.

For measurements with specific atmospheric O_2_ concentrations, a CLARIOstar® plate reader equipped with an atmospheric control unit (ACU) was used. By flushing N_2_ (99.998 % v/v, Westfalen AG, Münster, Germany) into the plate reader chamber, the ACU allows atmospheric oxygen concentrations to be set between 0.1 and 20% inside the plate reader chamber. For experiment at different O_2_ concentrations, the pump system of the plate reader was used to inject the solution. Pumps were primed with the flg22 solution prior to starting the measurement and an injection speed of 150 µL/s with smart dispensing was set. The elicitor or mock solutions were deoxygenated by N2 flushing before loading the solutions into the plate reader pumping system.

### Confocal imaging and ratiometric image analysis

Confocal microscopy was conducted at the LSM980 from Zeiss. The time series were performed with either 5x objective for x-y imaging or 25x objective for x-y-z imaging. Leaf discs from 28-35-day-old plants expressing the Peredox-mCherry or HyPer7 biosensor were dark-adapted for 2 hours and placed in one ibidi glass bottom µ-Dish (ibidi GmbH, Gräfelfing, Germany). Leaf discs were laterally fixed with a tape, which also served as a spacer to position a metal mesh over the sample to keep it in focus. Samples were imaged in 1mL plate reader assay medium following treatments of either 50 µL of buffer (mock) or flg22 (final concentration 1 µM).

For Peredox mCherry imaging, the tSapphire fluorophore was excited at 400 nm and emission was detected between 491 nm and 540 nm, while mCherry was excited at 561 nm and the emission measured between 588 nm and 632 nm. For HyPer7 imaging, the fluorophore was excited at 405 nm and 488 nm and emission detected between 508 and 535 nm. Imaging was done with an open pinhole. Chlorophyll autofluorescence was detected between 654 nm and 685 nm. For x–y–z imaging, a total of 287 optical sections were acquired at 3 µm intervals.

Time series were analyzed with the redox ratio analysis software RRA (https://markfricker.org/77-2/software/redox-ratio-analysis/) (*76*) and ratiometric videos prepared using Fiji (*77*). The ratios of the tSapphire and the mCherry channels (Peredox mCherry) and 405 nm and 488 nm channels (HyPer7) were analyzed based on the average ratio of each single picture. The minimal ratio was set to 0.1, the maximum ratio to 2.5. The used pixel intensities were set from 0 to 70. The background was set white.

### Respirometric assays

A total of 1 mL of plate reader assay buffer was used to reconstitute a vial containing dried MitoXpress® Xtra probe (Agilent Technologies, Santa Clara, CA, U.S.A) to create a stock solution. Single leaf discs excised from Arabidopsis plants, weighed and floated into each well of a transparent 96-well microplate containing 225µL of plate reader assay buffer and 25µL of MitoXpress Xtra stock solution. The plate was monitored in a CLARIOstar® plate reader pre-equilibrated at 25 °C in TR-F mode, measuring probe signal in each well every 30 s. Instrument TR-F settings were: excitation filter – TR, dichroic mirror – LP-TR, emission filter – 645-20 nm, Delay times of 30 µs and 70 µs (two windows). After 20 minutes, 100 µL of mineral oil pre-warmed at 37 °C was added to each well to seal the samples from ambient air. After an additional 20 minutes elicitor treatment was carried out by pipetting flg22 to a final concentration of 100nM below the mineral oil phase. Phosphorescence lifetime values (LT) for each sample and measurement point were calculated in Excel using the following formula according to the manufacturer’s protocol: LT= (t2-t1)/ln(F1/F2) where t1 and t2 are the delay times (30 and 70 µs) and F1 and F2 are the corresponding intensity signals. The resulting LT values for each assayed well were plotted and the slope of the oxygen concentration curve over time was used to calculate oxygen consumption rate for each sample. The fresh weight of each assayed leaf disc was used to normalize oxygen consumption rate values.

### GC-MS metabolite quantification

A total of 8 leaf discs were pooled from 4 different plants and transferred to 96-well plates containing 180uL of plate reader assay medium (10 mM MES, 10 mM MgCl_2_, 10 mM CaCl_2_, 5 mM KCl, pH 5.8 adjusted with KOH). Each pool of 8 leaf discs was considered one biological replicate. The plates were incubated in the dark for 2h 30 min before 20uL of either 1µM flg22 (100nM final concentration) or 20µL water (mock control) was added to each well. Leaf discs were collected 50 and 120 min after the treatment, dried using lint-free laboratory wipes (Kimwipes: Kimberly-Clark, Irving, TX, USA), and immediately flash-frozen in liquid nitrogen. Metabolite extraction was performed as previously described (*78*) with modifications. A total of 40 mg of frozen ground tissue was extracted in 400 µL pre-cooled (-20 °C) methanol containing 8 µg / mL ribitol (internal standard), followed by vortexing (1 min), shaking (15 min, 1000 rpm, 4 °C), and sonication (15 min). After addition of 400 µL chloroform and 400 µL water with intermediate vortexing, samples were centrifuged (14000 g, 5 min), and 200 µL of the upper phase was dried overnight. Derivatization was performed in 40 µL methoxyamine hydrochloride (20 mg / mL in pyridine, 120 min, 37 °C), followed by an addition of 70 µL N-Methyl-N-(trimethylsilyl)trifluoracetamide (30 min, 37 °C). Samples (1 µL) were injected in splitless mode using a Gerstel Multi-Purpose autosampler (Gerstel GmbH & Co.KG, Mülheim an der Ruhr, Germany) into a GC-TOF-MS system (Leco Pegasus HT TOF-MS; LECO Corporation, St. Joseph, MI, USA). Helium was used as carrier gas (2 mL s^-1^), and separation was performed on a DB-35 column (30 m x 0.32 mm x 0.25 µm). The oven temperature was increased from 85 °C to 360 °C at a rate of 15 °C min-1. Mass spectra were recorded after a 180 s solvent delay at 20 scans s-1 over a range of 70-600 m/z.

### Isotope labelling and GC-MS labelled metabolite quantification

A total of 8 leaf discs were pooled from 4 different plants, weighed, and transferred to 96-well plates containing 180uL of plate reader assay medium (10 mM MES, 10 mM MgCl_2_, 10 mM CaCl_2_, 5 mM KCl, pH 5.8 adjusted with KOH). Each pool of 8 leaf discs was considered one biological replicate. The plates were incubated in the dark for 2h 30 min before 20uL of either 1µM flg22 + 50mM 13C-U-Glucose (100nM flg22 + 5mM 13C-U-glucose final concentration) or 20µL of 50mM 13C-U-glucose (mock control) was added to each well. Leaf discs were collected 0, 60, 120 and 180 min after the treatment, dried using lint-free laboratory wipes (Kimwipes: Kimberly-Clark, Irving, TX, USA), and immediately flash-frozen in liquid nitrogen. Metabolite extraction was performed as previously described (*79*) with modifications. Eight leaf discs (approximately 40mg) were ground in a TissueLyzer (Qiagen, Hilden, Germany) for 1 min 30 s at 25 Hz. Subsequently, 500 µL extraction solution (methanol/chloroform/water 32.25:12.5:6.25, v/v/v) was added to each tube, followed by immediate vortexing. Samples were transferred to a cold-room at 4 °C and incubated under constant agitation for 30 min. Then, 500 µL of 0.005 mg/mL allo-inositol was added as an internal standard. Samples were centrifuged at 12,000 x *g* for 3 min, and the upper aqueous phase was transferred to a new 2 mL tube and flash-frozen in liquid N_2_. Samples were stored at -80 °C until analysis.

For analysis, 50µL of each extract was transferred to a new 2 mL tube and evaporated under a stream of N_2_. Derivatization was performed by adding 15 µL methoxyamine hydrochloride (30 mg / mL in pyridine anhydrous) with overnight incubation at room temperature, followed by the addition of 30 µL N-Methyl-N-(trimethylsilyl)trifluoracetamide (incubated for at least 1 h). Samples (1 µL) were injected in splitless and split modes using a PAL3 autosampler (CTC Analytics AG, Switzerland) into a GC-TOF-MS system (8890 GC System and 7250 GC/Q-TOF, Agilent Technologies, Inc., Santa Clara, United States). The inlet temperature was set to 250 °C. Helium was used as carrier gas (1.1 mL/min), and separation was performed on a HP-5ms Ultra Inert column (30 m x 250 μm x 0.25 μm, Agilent 19091S-433UI). The oven temperature was increased from 50 °C to 320 °C at a rate of 10 °C min-1. Mass spectra were recorded after a 6.5 min solvent delay over a range of 50-500 m/z.

### Statistical analysis

All statistical analyses were performed using GraphPad Prism 8.0.1 (GraphPad Software, Boston, MA, USA). Data were tested for normality prior to statistical analysis, and data that did not meet the assumptions of parametric tests were log_10_-transformed where appropriate. For comparisons involving a single independent variable, one-way analysis of variance (ANOVA) followed by Tukey’s multiple-comparisons test was used. For experiments involving two independent variables, two-way ANOVA followed by Tukey’s multiple-comparisons test was performed. A p-value < 0.05 was considered statistically significant. For metabolomics, Tukey-adjusted pairwise comparisons were additionally corrected across metabolites using the Benjamini–Hochberg procedure. An adjusted p-value (q-value) < 0.05 was considered statistically significant. The statistical test used for each experiment, as well as the number of biological replicates, is indicated in the respective figure legends.

## Supplemental Figures

**Figure S1.**
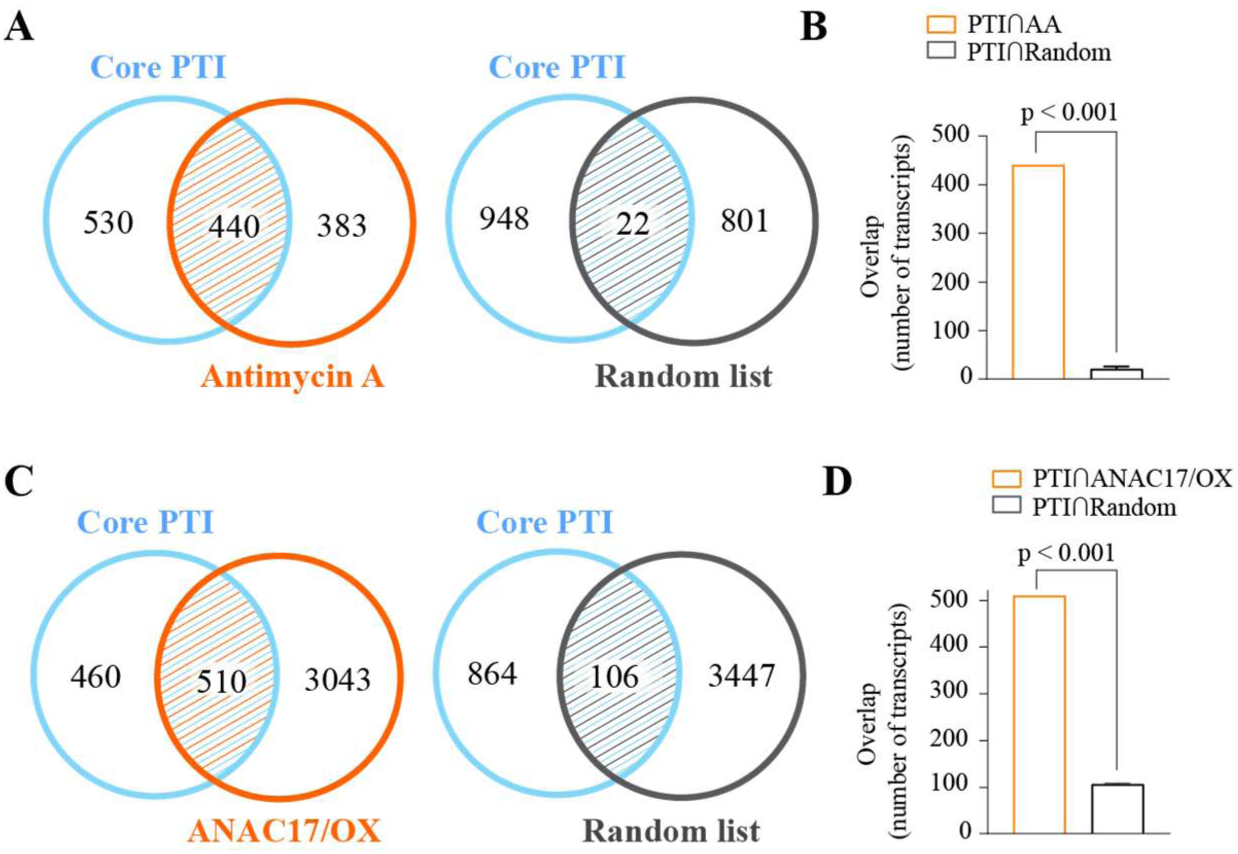
Overlap of core PTI-responsive transcripts with antimycin A- and ANAC17-overexpression-responsive genes. Overlap between core PTI-responsive transcripts and transcripts responsive to **(A, B)** antimycin A, and **(C, D)** ANAC17 overexpression (ANAC17/OX). ANAC17 overexpression-responsive transcripts were obtained from Meng et al. (2019) (P < 0.05, log₂FC > 1). Antimycin A-responsive transcripts were obtained from Zhu et al. (2023) (P < 0.05, log₂FC > 1). Core PTI-responsive transcripts were defined according to Bjornson et al. (2021) as transcripts significantly induced (P < 0.05, log₂FC > 1) by all tested PTI elicitors. **(A, C)** Venn diagrams show the observed overlap between each transcript set and the core PTI-responsive transcript set. **(B, D)** Bar graphs compare the observed overlap with the expected overlap, estimated as the mean overlap obtained from three randomly generated gene lists of equal size sampled from the TAIR12 genome annotation. Statistical significance of the enrichment was assessed using two-sided chi-square tests.

**Figure S2.**
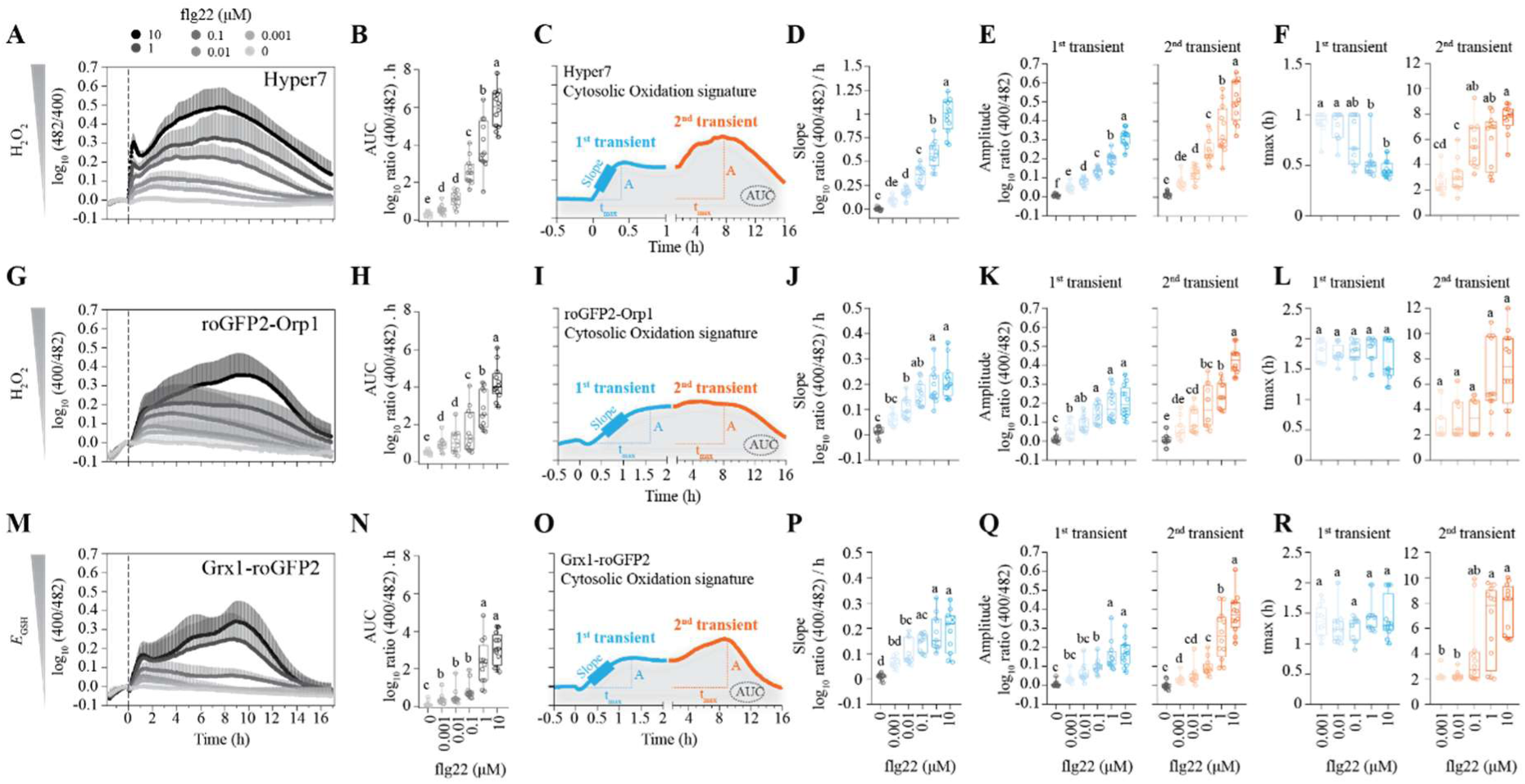
Quantitative analysis of biphasic H₂O₂ and *E*_GSH_ dynamics following flg22 treatment. In vivo flg22 dose-response analyses were performed using **(A-F)** HyPer7, **(G-L)** roGFP2-Orp1, and **(M–R)** Grx1-roGFP2 biosensor lines. **(A, G, M)** In vivo biosensor oxidation dynamics following treatment with increasing concentrations of flg22. Dashed lines indicate flg22 treatment. Lines represent mean values. Error bars represent SD. **(B, H, N)** Area under the curve (AUC) of the corresponding oxidation traces. **(C, I, O)** Schematic illustrating the parameters extracted from the biosensor responses, including the initial oxidation rate, maximal amplitude, and time to reach maximal amplitude. **(D, J, P)** Initial oxidation rate. **(E, K, Q)** Maximal oxidation amplitude. **(F, L, R)** Time to reach maximal oxidation amplitude. Error bars shown on boxplots represent minimum to maximum values. n = 12 biological replicates for all panels. Different letters indicate statistically significant differences among treatments (P < 0.05, one-way ANOVA followed by Tukey’s multiple-comparison test).

**Figure S3.**
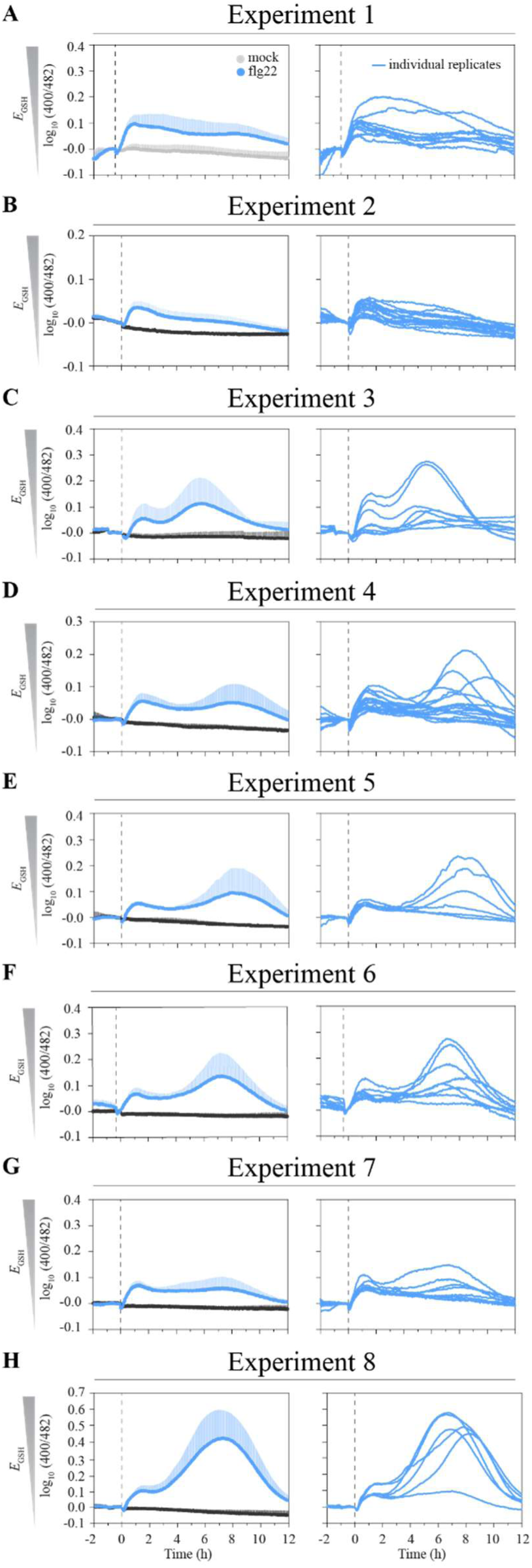
Experimental variability of flg22-induced *E*_GSH_ responses. (A–H) In vivo oxidation dynamics of the Grx1-roGFP2 biosensor following treatment with 100 nM flg22 across independent experiments. Dashed lines indicate the time of flg22 application. Lines on left panels are mean values. Error bars represent SD. Lines on right panels show the respective individual biological replicates from each independent experiment shown on the left.

**Figure S4.**
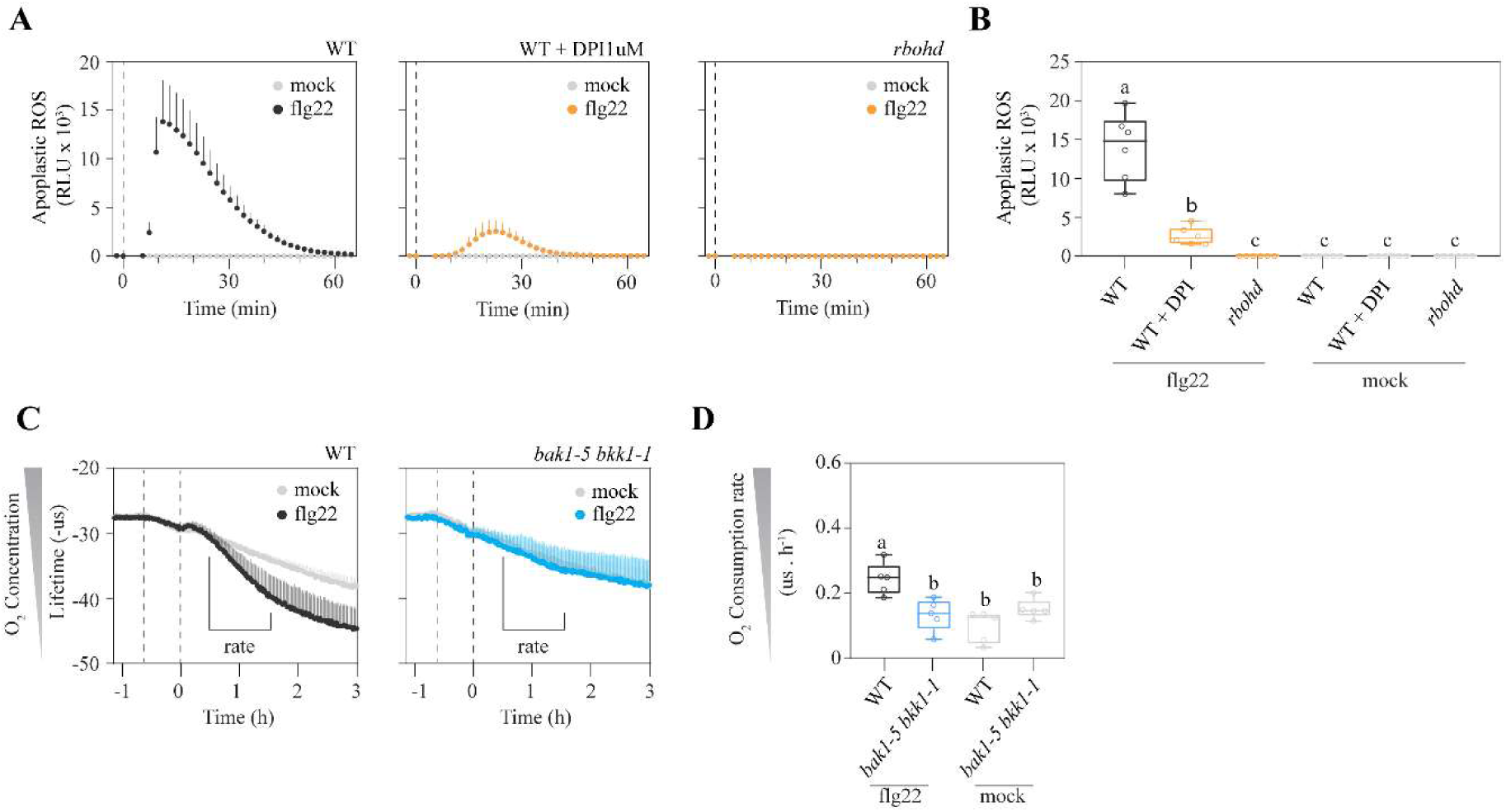
Apoplastic ROS burst and respiratory responses in different genetic backgrounds. **(A)** Apoplastic ROS burst in RBOHD mutant and DPI-treated plants following treatment with 100 nM flg22. Dashed lines indicate the time of flg22 application. n = 6 biological replicates. Error bars represent SD. **(B)** Quantification of the amplitude of the elicitor-induced apoplastic ROS burst derived from (A). Error bars represent minimum to maximum values. **(C)** Oxygen consumption measured using the MitoXPress assay in WT and *bak1-5 bkk1-1* mutants. The first dashed line indicates the addition of mineral oil, and the second dashed line (time = 0 h) indicates treatment with 100 nM flg22. n = 5 biological replicates. Error bars represent SD. **(D)** Oxygen consumption rate derived from (C), calculated as the slope between 0.5 and 1.5 h after flg22 treatment. Error bars represent minimum to maximum values. Different letters indicate statistically significant differences among genotypes (*P* < 0.05, one-way ANOVA followed by Tukey’s multiple-comparison test).

**Figure S5.**
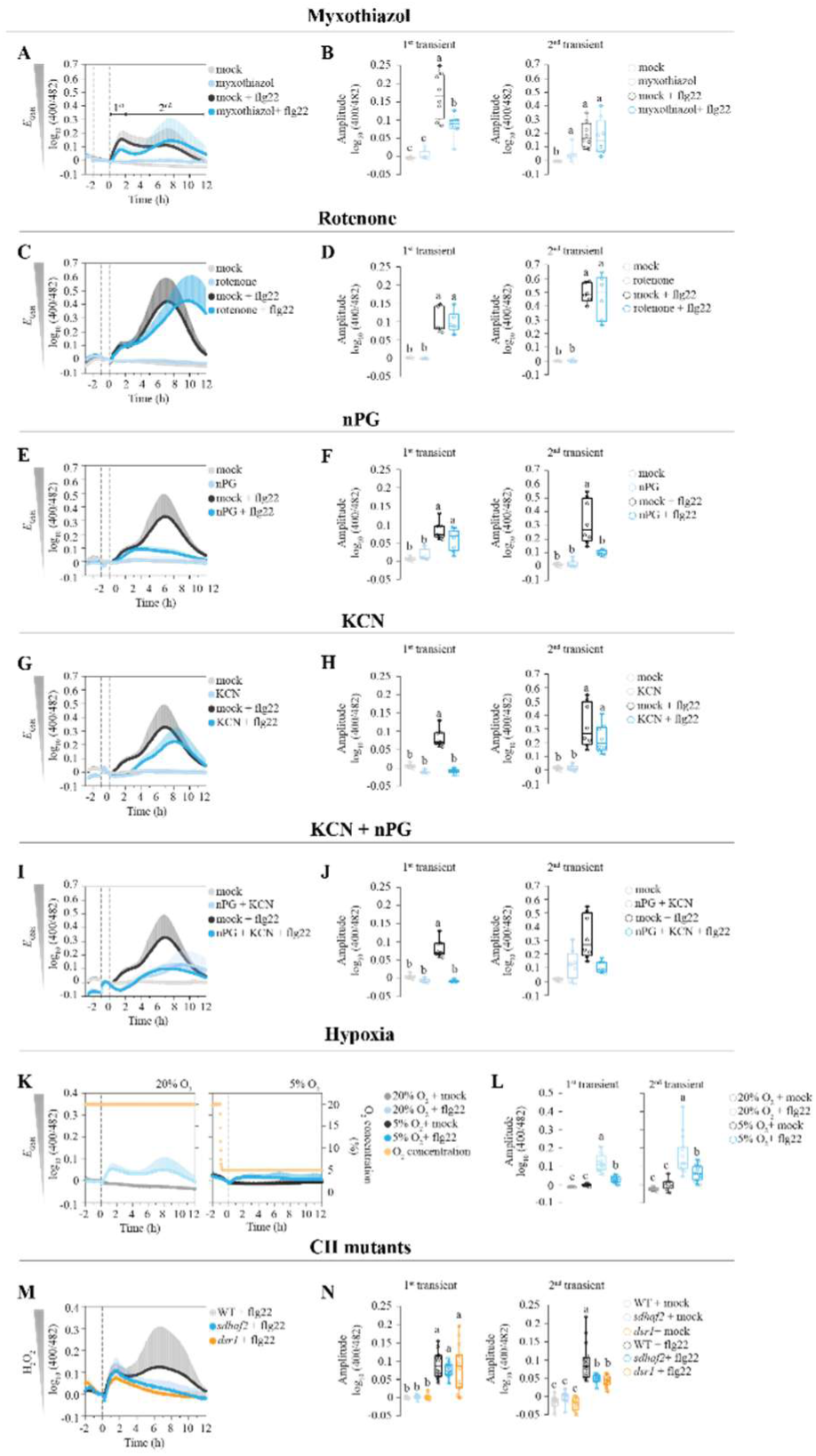
Pharmacological, environmental, and genetic perturbations alter elicitor-induced *E*_GSH_ responses. In vivo oxidation dynamics of the Grx1-roGFP2 biosensor in Arabidopsis thaliana leaf discs following treatment with 100 nM flg22 after pre-treatment with (**A, B)** myxothiazol (50 μM, n = 8 biological replicates), **(C, D)** rotenone (50 μM, n = 6 biological replicates), **(E, F)** n-propyl gallate (nPG, 500 μM, n = 8 biological replicates), **(G, H)** KCN (1 mM, n = 8 biological replicates). **(I, J)** KCN (1 mM) + nPG (500 μM, n = 8 biological replicates), or **(K, L)** low oxygen (5% O₂, n = 16 biological replicates). **(M, N)** In vivo oxidation dynamics of the roGFP2-Orp1 biosensor in Arabidopsis thaliana leaf discs following treatment with 100 nM flg22 in the complex II mutants *dsr1* and *sdhaf2*. n = 16 biological replicates. Left panels show *E*_GSH_ time courses following elicitor treatment. Lines represent mean values. Error bars represent SD. The first dashed line indicates pharmacological treatments. The second dashed line (time = 0h) indicates 100nM flg22 treatment. Right panels show the amplitudes of the first and second transients, where the first transient was defined as the maximum response between 0–2 h and the second transient as the maximum response between 2–12 h. Error bars represent minimum to maximum values. **(A–D)** 0.1% ethanol was used as a mock control. Water was used as a mock treatment for the other panels. **(E–J)** Panels are from the same experiment plotted in different panels. Different letters indicate statistically significant differences among treatments or genotypes (one-way ANOVA followed by Tukey’s multiple-comparison test, P < 0.05).

**Figure S6.**
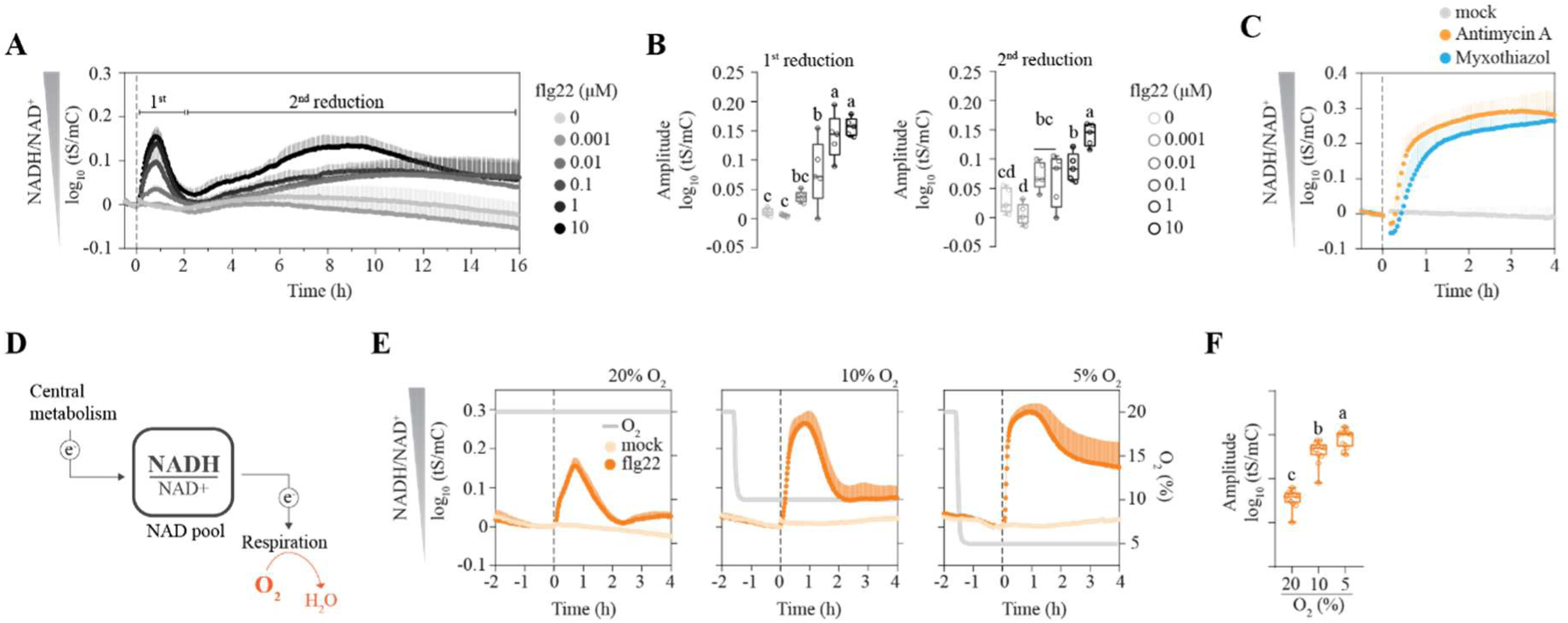
Quantitative analysis of cytosolic NAD redox dynamics following flg22 treatment. **(A)** Time courses of cytosolic NADH/NAD⁺ dynamics following flg22 treatment. Lines represent mean values. n = 5 biological replicates. Error bars represent SD. **(B)** Quantification of the amplitudes of the first and second NAD reduction phases in response to increasing flg22 concentrations. The first reduction phase was defined as the maximum increase in the NADH/NAD⁺ ratio between 0 and 2 h after treatment, whereas the second reduction phase was defined as the maximum response between 2 and 12 h. Error bars represent minimum to maximum values. **(C)** Cytosolic NADH/NAD⁺ dynamics following treatment with the mitochondrial electron transport chain inhibitors antimycin A (50 μM) and myxothiazol (50 μM). Dashed line (time = 0h) indicates the inhibitor application. Plants treated with 0.1% ethanol served as the mock control. **(D)** Schematic illustration of the major sources and sinks of electrons within the cellular NAD pool. **(E)** Time courses of cyto-nuclear NADH/NAD⁺ dynamics following flg22 treatment in plants maintained under 20%, 10%, or 5% O₂. Lines represent mean values. Error bars represent SD. **(F)** Quantification of the amplitude of the first NAD reduction phase under different oxygen concentrations. Error bars represent SD. Different letters indicate statistically significant differences among treatments or genotypes, as determined by one-way ANOVA followed by Tukey’s multiple-comparison test (P < 0.05).

**Figure S7.**
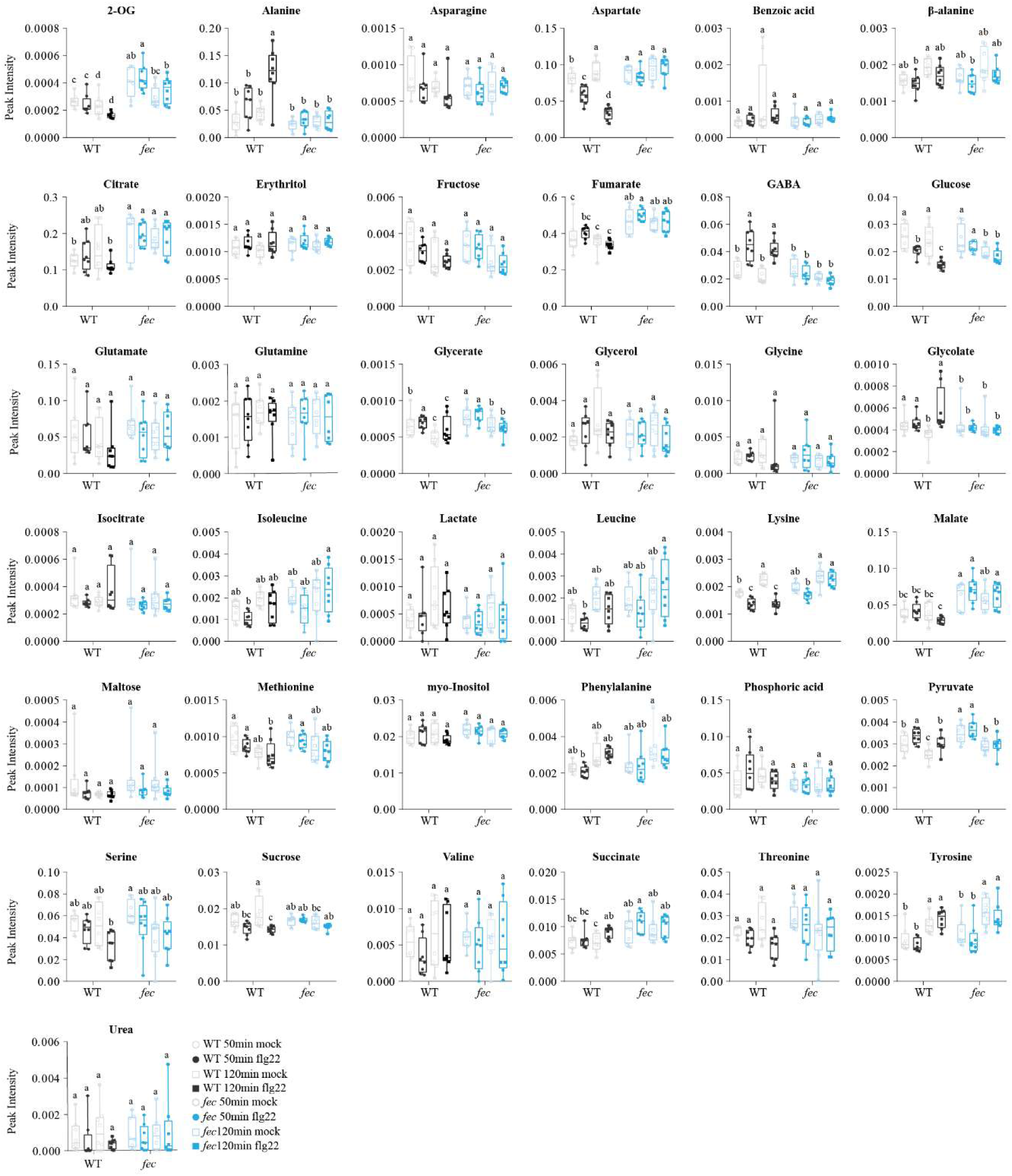
Untargeted GC-MS metabolite quantification in elicitor-treated WT and *fec* plants. Different letters indicate statistically significant differences among treatments or genotypes, as determined by two-way ANOVA followed by Tukey’s multiple-comparisons test. p-values were adjusted for multiple testing using the Benjamini–Hochberg false discovery rate (FDR) correction. An adjusted p-value (q-value) < 0.05 was considered statistically significant. Error bars represent minimum to maximum values. n = 8 biological replicates.

**Figure S8.**
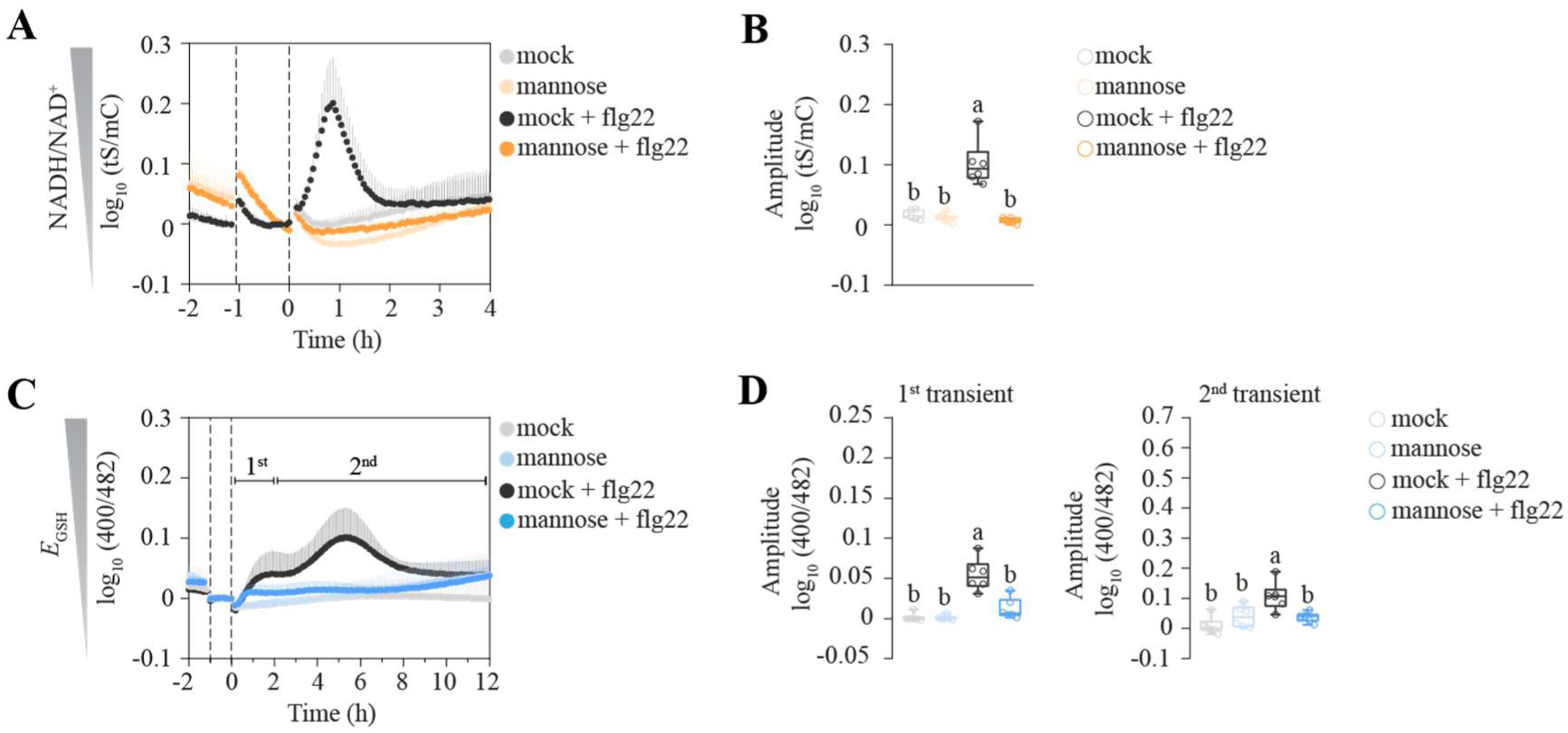
Elicitor-induced NAD redox and *E*_GSH_ dynamics in response to mannose. **(A)** Time course of cytosolic NADH/NAD^+^ dynamics following combined 10mM mannose and 100nM flg22 treatment. Lines represent mean values. Error bars represent SD. **(B)** Quantification of the amplitude of the first elicitor-induced NAD reduction phase in the presence of mannose. The amplitude was defined as the maximum increase in the NADH/NAD⁺ ratio between 0 and 2 h after treatment. Error bars represent minimum to maximum values. **(C)** Time course of *E*_GSH_ dynamics following combined 10mM mannose and 100nM flg22 treatment. **(D)** Quantification of the amplitudes of the first and second elicitor-induced *E*_GSH_ oxidation phases in the presence of mannose. The first transient oxidation was defined as the maximum oxidation response of *E*_GSH_ between 0 and 2 h after treatment, whereas the second oxidation phase was defined as the maximum response between 2 and 12 h. Different letters indicate statistically significant differences among treatments, as determined by one-way ANOVA followed by Tukey’s multiple-comparison test (*P* < 0.05).

**Figure S9.**
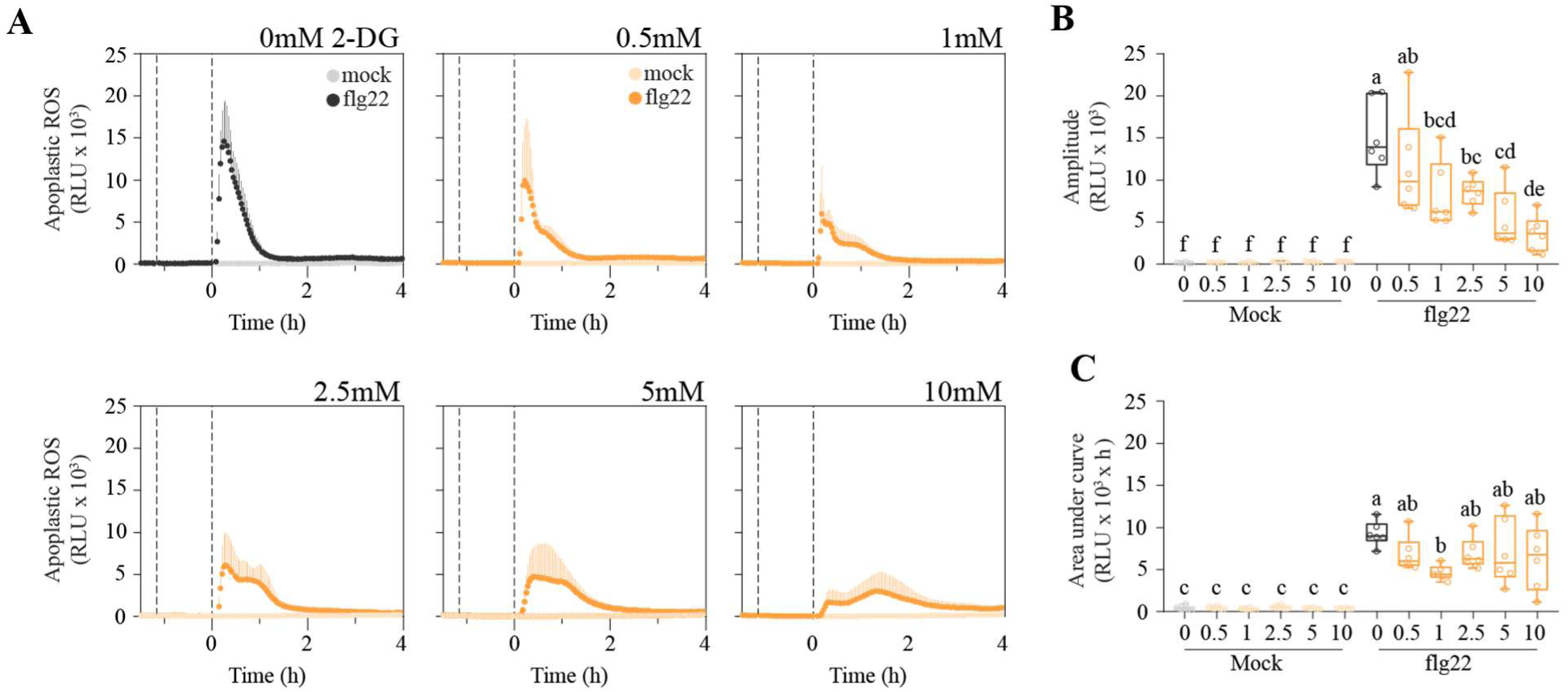
Elicitor-induced apoplastic ROS burst in response to 2-deoxyglucose (2-DG). **(A)** Apoplastic ROS burst after 100nM flg22 treatment in Arabidopsis thaliana leaf discs pre-treated with different concentrations of 2-DG. The first dashed line indicates 2-DG treatment. The second dashed line (time = 0h) indicates flg22 treatment. Lines represent mean values. Error bars represent SD. **(B)** Quantification of the amplitude of the elicitor-induced apoplastic ROS burst in the presence of 2-DG. **(C)** Quantification of area under the curve of the elicitor-induced apoplastic ROS burst in the presence of 2-DG. (B, C) Error bars represent minimum to maximum values. n = 6 biological replicates for all panels. Different letters indicate statistically significant differences among treatments, as determined by one-way ANOVA followed by Tukey’s multiple-comparison test (*P* < 0.05).

**Figure S10.**
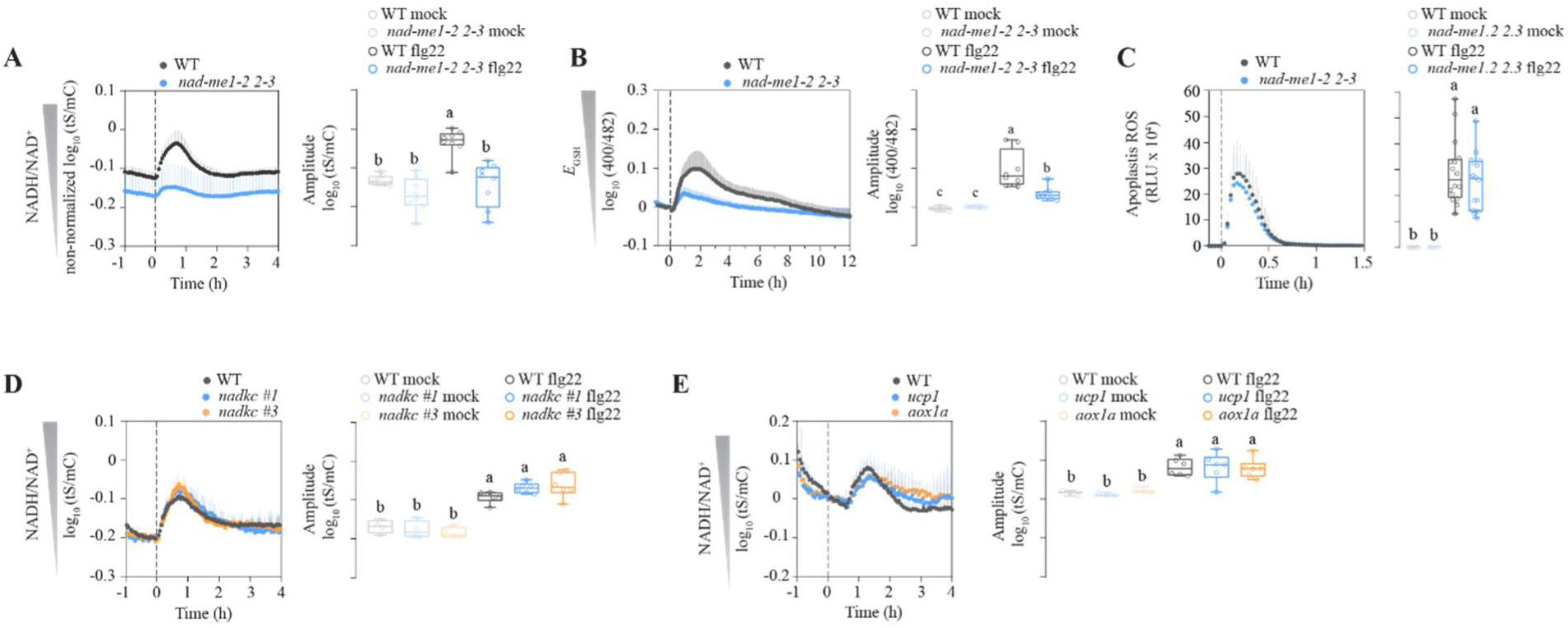
Elicitor-induced NAD redox, *E*_GSH_ dynamics and apoplastic ROS burst in malate metabolism, *nadkc*, *ucp1* and *aox1a* mutants. **(A)** Time course of NADH/NAD^+^ dynamics following flg22 treatment in *nad-me1-2 2-3* double mutant (left). Lines represent mean values. Error bars represent SD. Quantification of the amplitude of the first elicitor-induced NAD reduction phase in the *nad-me1-2 2-3* double mutant (right). Error bars represent minimum to maximum values. n = 8 biological replicates. **(B)** Time course of *E*_GSH_ dynamics following flg22 treatment in *nad-me1-2 2-3* double mutant (left). Lines represent mean values. Error bars represent SD. Quantification of the amplitude of the first elicitor-induced *E*_GSH_ oxidation phase in the *nad-me1-2 2-3* double mutant (right). Error bars represent minimum to maximum values. n = 8 biological replicates. **(C)** Apoplastic ROS burst after 100nM flg22 treatment in *nad-me1-2 2-3* double mutant (left). Lines represent mean values. Error bars represent SD. Quantification of the amplitude of the elicitor-induced apoplastic ROS burst in the *nad-me1-2 2-3* double mutant (right). Error bars represent minimum to maximum values. n = 16 biological replicates. **(D)** Time course of NADH/NAD^+^ dynamics following 100nM flg22 treatment in *nadkc* mutant (left). Lines represent mean values. Error bars represent SD. Quantification of the amplitude of the first elicitor-induced NAD reduction phase in the *nadkc* mutant (right). *nadkc#1 and #3* are two independent transgenic lines carrying the biosensor Peredox mCherry. Error bars represent minimum to maximum values. n = 6 biological replicates. **(E)** Time course of NADH/NAD^+^ dynamics following 100nM flg22 treatment in *ucp1* and *aox1a* single mutants (left). Lines represent mean values. Error bars represent SD. Quantification of the amplitude of the first elicitor-induced NAD reduction phase in the *ucp1* and *aox1a* single mutants (right). Error bars represent minimum to maximum values. n = 6 biological replicates. Dashed line (time = 0h) indicates flg22 treatment. Different letters indicate statistically significant differences among treatments, as determined by one-way ANOVA followed by Tukey’s multiple-comparison test (*P* < 0.05).

**Figure S11.**
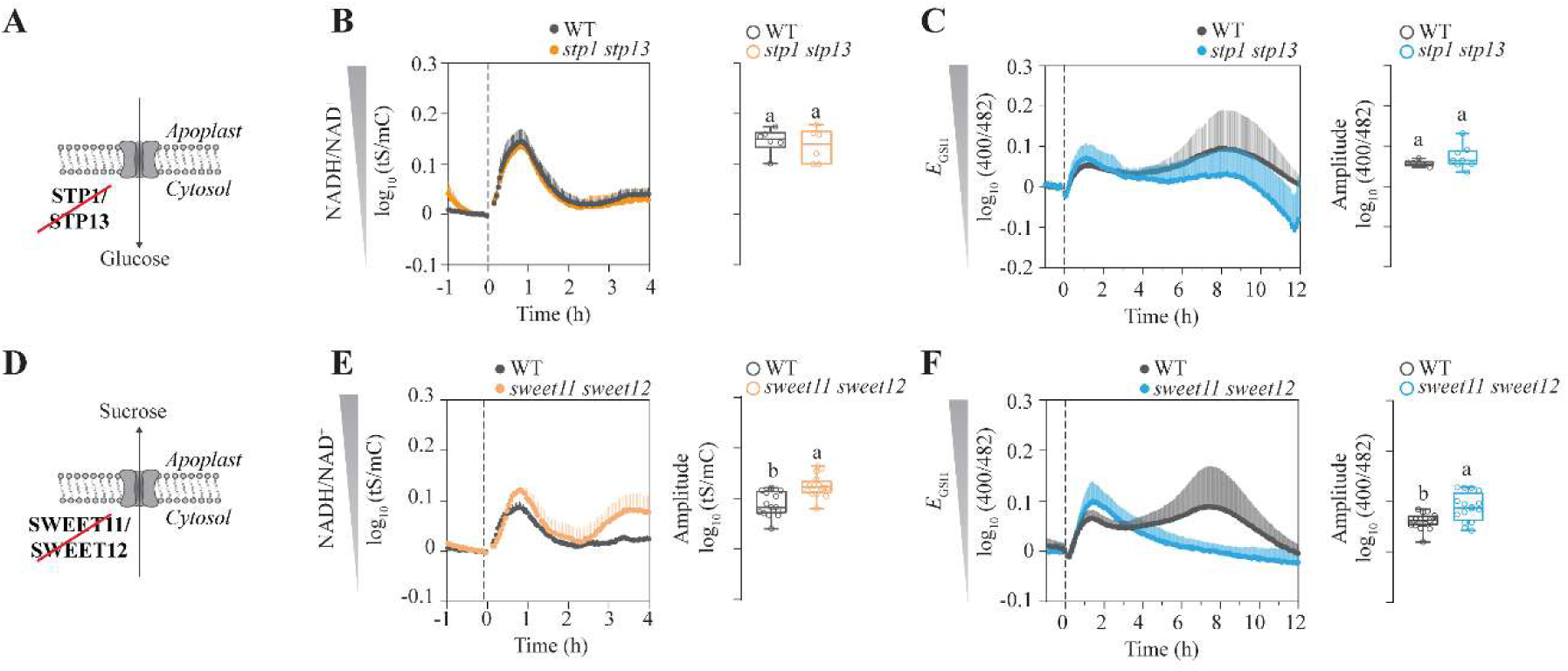
Elicitor-induced NAD redox and *E*_GSH_ dynamics in sugar transport or breakdown mutants. **(A)** Schematic representation of STP1 and STP13 function. **(B)** Time course of NADH/NAD^+^ dynamics following 100 nM flg22 treatment in *stp1 stp13* double mutant (left). Quantification of the amplitude of the first elicitor-induced NAD reduction phase in the *stp1 stp13* double mutant (right). **(C)** Time course of *E*_GSH_ dynamics following 100 nM flg22 treatment in *stp1 stp13* double mutant (left). Quantification of the amplitude of the first elicitor-induced *E*_GSH_ oxidation phase in the *stp1 stp13* double mutant (right). **(D)** Schematic representation of SWEET11 and SWEET12 function. **(E)** Time course of NADH/NAD^+^ dynamics following 100 nM flg22 treatment in *sweet11 sweet12* double mutant (left). Quantification of the amplitude of the first elicitor-induced NAD reduction phase in the *sweet11 sweet12* double mutant (right). **(F)** Time course of *E*_GSH_ dynamics following 100 nM flg22 treatment in *sweet11 sweet12* double mutant (left). Quantification of the amplitude of the first elicitor-induced *E*_GSH_ oxidation phase in the *sweet11 sweet12* double mutant (right). Dashed lines (time = 0h) indicate flg22 treatment. Water was used as a mock control for all panels. Different letters indicate statistically significant differences among treatments, as determined by one-way ANOVA followed by Tukey’s multiple-comparison test (*P* < 0.05).

**Figure S12.**
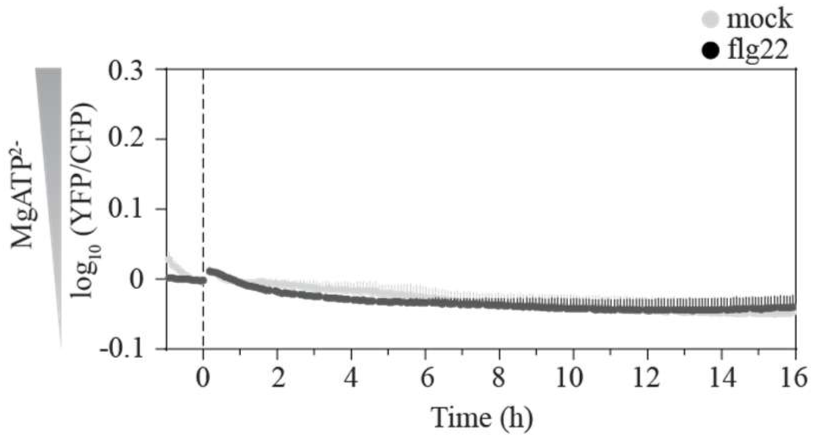
Time courses of cytosolic MgATP^2-^dynamics following flg22 treatment. Lines represent mean values. n = 8 biological replicates. Error bars represent SD. Dashed line indicates 100 nM flg22 treatment. Water was used as a mock control.

**Movie S1.**

Representative confocal time-course showing H₂O₂ dynamics in the leaf epidermis following 1 µM flg22 treatment at the indicated time points, monitored using HyPer7. Scale bars = 100 µm. (provided as a separate file)

**Movie S2.**

Representative confocal time-course of H₂O₂ dynamics following 1 µM flg22 treatment at 0, 0.25, 6, and 15 h. H₂O₂ levels were monitored using HyPer7. Scale bar = 10 µm. (provided as a separate file)

Table S1. Biological processes enriched in the 1,000 PTI-upregulated genes (provided as a separate file).

Table S2. Mitochondrial dysfunction regulon transcript responses after treatment with different elicitors (provided as a separate file).

Table S3. Intersection between the core PTI-upregulated genes and either antimycin A or ANAC17/OE upregulated genes (provided as a separate file).

Table S4. Untargeted GC-MS metabolite analysis of WT and *fec* plants treated with 100 nM flg22 for 50 and 120 min (provided as a separate file).

**Table S5.**
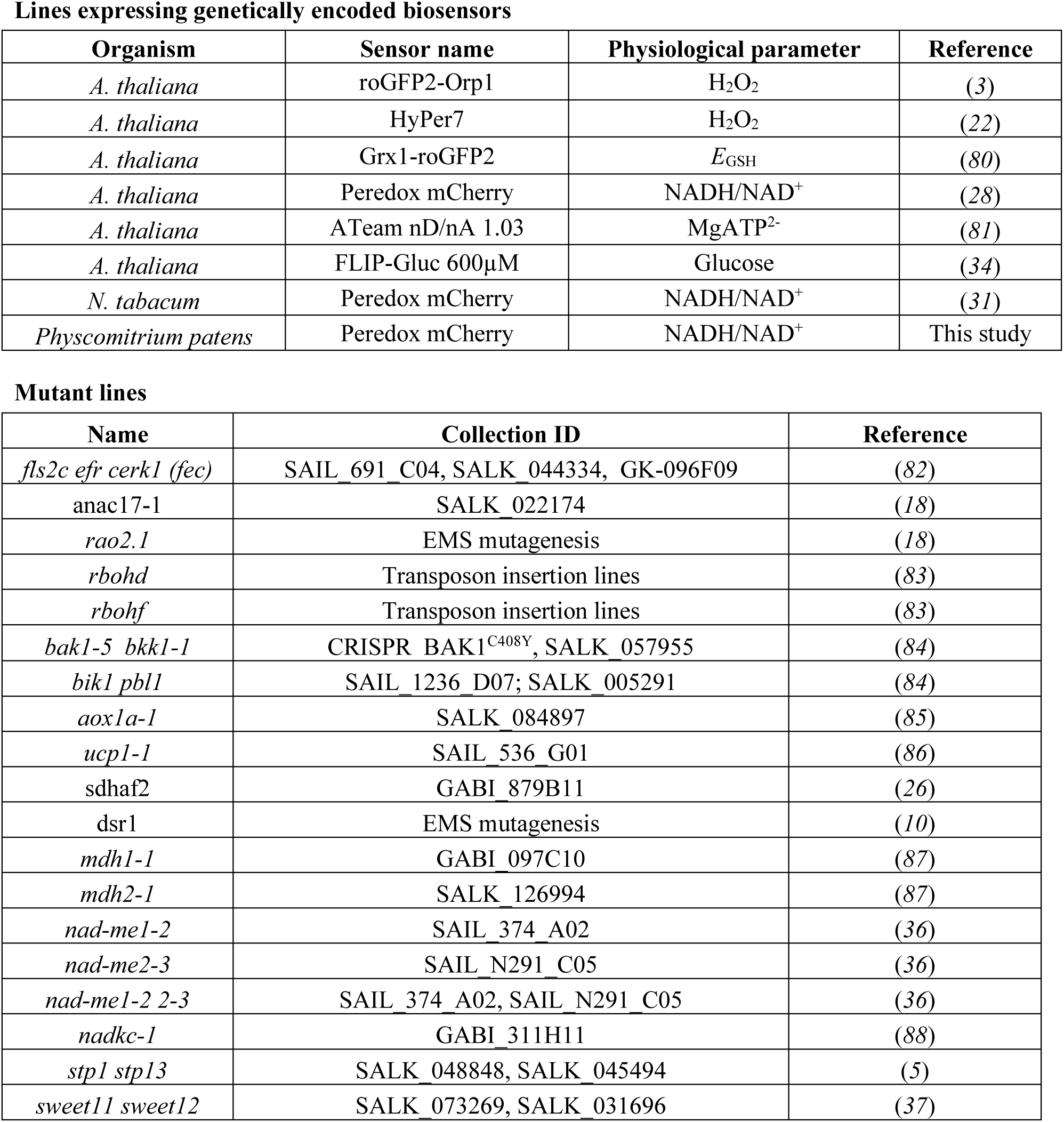
List of plant material used in this study.

| **Lines expressing genetically encoded biosensors** |  |  |  |
| **Organism** | **Sensor name** | **Physiological parameter** | **Reference** |
|---|---|---|---|
| *A. thaliana* | roGFP2-Orp1 | H<sub>2</sub>O<sub>2</sub> | (*3*) |
| *A. thaliana* | HyPer7 | H<sub>2</sub>O<sub>2</sub> | (*22*) |
| *A. thaliana* | Grx1-roGFP2 | *E*<sub>GSH</sub> | (*80*) |
| *A. thaliana* | Peredox mCherry | NADH/NAD<sup>+</sup> | (*28*) |
| *A. thaliana* | ATeam nD/nA 1.03 | MgATP<sup>2-</sup> | (*81*) |
| *A. thaliana* | FLIP-Gluc 600µM | Glucose | (*34*) |
| *N. tabacum* | Peredox mCherry | NADH/NAD<sup>+</sup> | (*31*) |
| *Physcomitrium patens* | Peredox mCherry | NADH/NAD<sup>+</sup> | This study |
| **Mutant lines** |  |  |  |
| **Name** | **Collection ID** | **Reference** |  |
| *fls2c efr cerk1 (fec)* | SAIL_691_C04, SALK_044334, GK-096F09 | (*82*) |  |
| anac17-1 | SALK_022174 | (*18*) |  |
| *rao2.1* | EMS mutagenesis | (*18*) |  |
| *rbohd* | Transposon insertion lines | (*83*) |  |
| *rbohf* | Transposon insertion lines | (*83*) |  |
| *bak1-5 bkk1-1* | CRISPR BAK1<sup>C408Y</sup>, SALK_057955 | (*84*) |  |
| *bik1 pbl1* | SAIL_1236_D07; SALK_005291 | (*84*) |  |
| *aox1a-1* | SALK_084897 | (*85*) |  |
| *ucp1-1* | SAIL_536_G01 | (*86*) |  |
| sdhaf2 | GABI_879B11 | (*26*) |  |
| dsr1 | EMS mutagenesis | (*10*) |  |
| *mdh1-1* | GABI_097C10 | (*87*) |  |
| *mdh2-1* | SALK_126994 | (*87*) |  |
| *nad-me1-2* | SAIL_374_A02 | (*36*) |  |
| *nad-me2-3* | SAIL_N291_C05 | (*36*) |  |
| *nad-me1-2 2-3* | SAIL_374_A02, SAIL_N291_C05 | (*36*) |  |
| *nadkc-1* | GABI_311H11 | (*88*) |  |
| *stp1 stp13* | SALK_048848, SALK_045494 | (*5*) |  |
| *sweet11 sweet12* | SALK_073269, SALK_031696 | (*37*) |  |

**Table S6.**
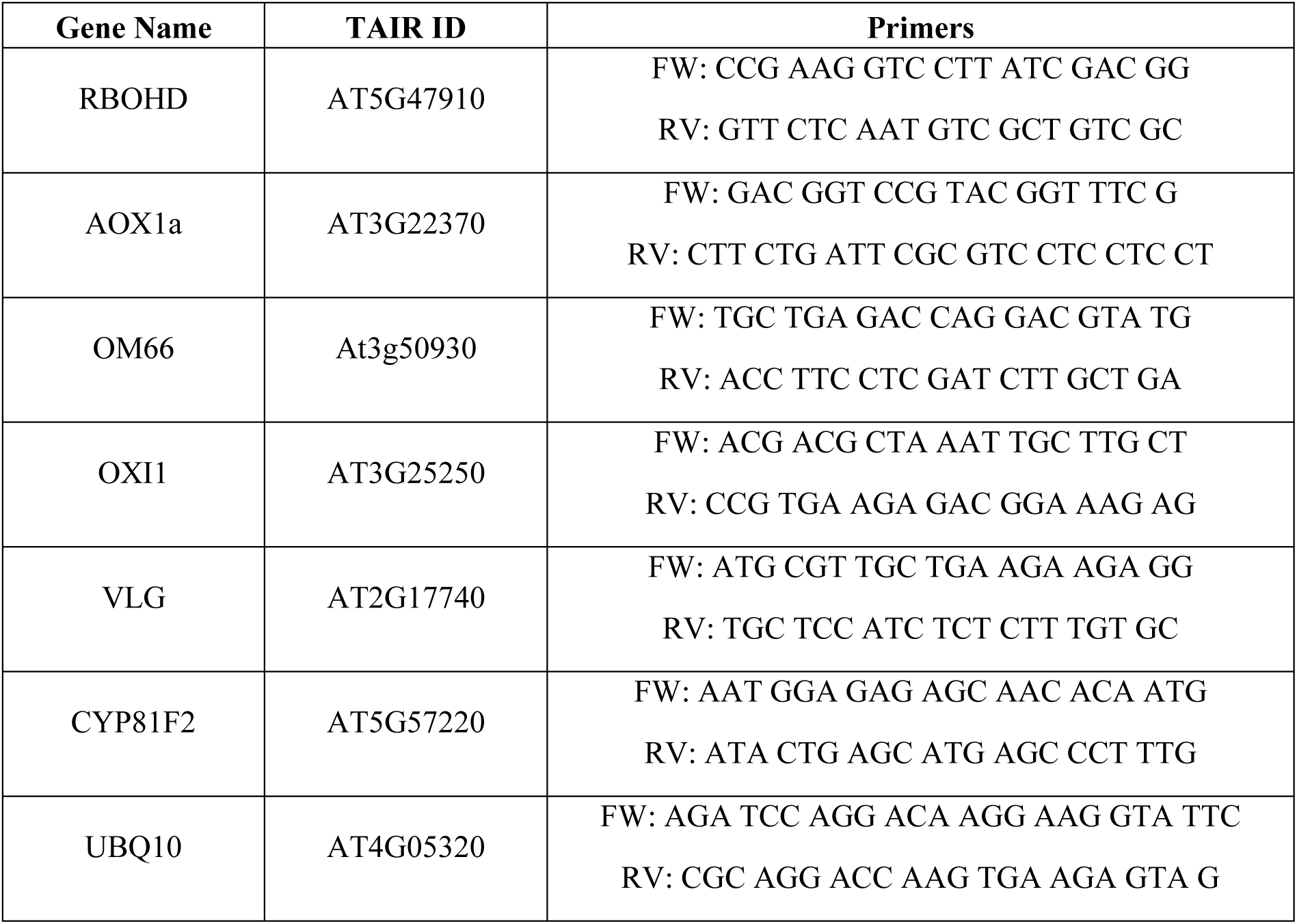
List of primers used in this study.

## Notes

### Competing Interest Statement

The authors have declared no competing interest.

